# Endometrial epithelial ARID1A is critical for uterine gland function in early pregnancy establishment

**DOI:** 10.1101/2020.09.23.308528

**Authors:** Ryan M. Marquardt, Tae Hoon Kim, Jung-Yoon Yoo, Hanna E. Teasley, Asgerally T. Fazleabas, Steven L. Young, Bruce A. Lessey, Ripla Arora, Jae-Wook Jeong

## Abstract

Though endometriosis and infertility are clearly associated, the pathophysiological mechanism remains unclear. Previous work has linked endometrial ARID1A loss to endometriosis-related endometrial non-receptivity. Here, we show in mice that ARID1A binds and regulates transcription of the *Foxa2* gene required for endometrial gland function. Uterine specific deletion of *Arid1a* compromises gland development and diminishes *Foxa2* and *Lif* expression. Deletion of *Arid1a* with Ltf-iCre in the adult mouse endometrial epithelium preserves gland development while still compromising gland function. Mice lacking endometrial epithelial *Arid1a* are severely sub-fertile due to defects in implantation, decidualization, and endometrial receptivity from disruption of the LIF-STAT3-EGR1 pathway. FOXA2 is also reduced in the endometrium of women with endometriosis in correlation with diminished ARID1A, and both ARID1A and FOXA2 are reduced in non-human primates induced with endometriosis. Our findings describe a role for ARID1A in the endometrial epithelium supporting early pregnancy establishment through the maintenance of gland function.

## Introduction

A primary function of the uterus is to support fertility by protecting and nourishing an embryo as it develops into a fetus and matures until birth. The inner layer of the uterus, the endometrium, is composed of a luminal epithelial cell layer surrounded by a supportive stromal cell layer containing epithelial gland structures. In the presence of an embryo, these distinct cell types coordinate through complex epithelial-stromal crosstalk to facilitate implantation and the establishment of a healthy pregnancy (1).

The ovarian steroid hormones progesterone (P4) and estrogen (E2) govern the human menstrual cycle in a 28-30 day process where E2-driven proliferation builds endometrial thickness in the proliferative phase before giving way to the P4-dominated secretory phase, which contains the transient window of embryo receptivity that depends on the length of P4 exposure (2). In order for successful pregnancy establishment to take place, endometrial epithelial cells must cease proliferation to allow embryo invasion, and stromal cells must be ready to differentiate into epithelioid secretory cells in a process called decidualization (3). In mice a parallel process occurs, but instead of a menstrual cycle, mice undergo a 4-5 day estrous cycle, in which the implantation window opens with a nidatory E2 surge (3, 4).

P4 and E2 maintain a tightly regulated, dynamic balance in the endometrium as they enact downstream signaling pathways, primarily through their cognate receptors, the progesterone receptor (PGR) isoforms PR-A and PR-B, and the estrogen receptors (ESR1 and ESR2) (3, 4). However, dysregulation of P4 and E2 signaling is common in uterine diseases such as endometriosis (5). Endometriosis occurs when endometrium-like tissue grows outside the uterus. Affecting about 1 in 10 women of reproductive age, this common disease frequently causes dysmenorrhea, chronic pelvic pain, and loss of fertility (6). Severe endometriosis can compromise fertility by directly diminishing ovarian reserve through endometriomas or by the distorting of pelvic anatomy, but these mechanisms do not explain the fertility defects observed in mild cases of endometriosis when endometrial receptivity is apparently affected (7, 8).

The murine nidatory E2 surge also induces endometrial gland secretion of leukemia inhibitory factor (LIF), a cytokine necessary for implantation and decidualization (3). Abundant evidence in several mammalian species supports the essential role of uterine glands and secretion of LIF in processes necessary for pregnancy success including implantation, decidualization, and placentation (9, 10). LIF expression is reportedly decreased in the glandular epithelium of women with endometriosis (11), which may reflect more general gland dysfunction. Forkhead box a2 (FOXA2), one of three FoxA family transcription factors involved in the development and function of many organs (12), is necessary for endometrial gland development, LIF expression, and pregnancy establishment in mice (10, 13). Being the only FoxA family member expressed in the mouse and human uterus, FOXA2 is specific to the glandular epithelium, and its mutation or loss of expression has been reported in uterine diseases such as endometrial cancers (14) and endometriosis (15–17). Though its physiological function in the human endometrium is not well characterized, recent evidence indicates that FOXA2 coordinates with other factors and pathways critically involved in uterine receptivity, implantation, and decidualization (18).

AT-rich interaction domain 1A (ARID1A), a 250 kDa switch/sucrose non-fermentable (SWI/SNF) chromatin remodeling complex subunit with known tumor suppressor function, has been linked to both endometriosis and regulation of endometrial receptivity (19–21). Though it is commonly mutated in endometriosis-associated ovarian cancers (22), the role of ARID1A in normal uterine physiology and in benign diseases such as endometriosis is not well understood. In normal conditions, ARID1A maintains strong nuclear expression in all uterine compartments throughout the menstrual cycle in women and throughout early pregnancy in mice (20). However, inactivating mutations in the *ARID1A* gene have been identified in deeply infiltrating endometriotic lesions (21) and ovarian endometriomas (23), and ARID1A expression is reduced in eutopic endometrial epithelium and stroma from women with endometriosis (20). Though other cancer-associated genes are frequently mutated in the normal eutopic endometrium and in that of women of endometriosis, *ARID1A* mutations are very rare (23, 24), implying that the decrease in ARID1A expression in the endometrium of women with endometriosis likely takes place at the epigenetic, transcriptional, or post-transcriptional level. In mice, deletion of uterine *Arid1a* causes infertility due to implantation and decidualization defects, increased E2-induced epithelial proliferation, and decreased epithelial P4 signaling at pre-implantation (20). Based on these findings, we hypothesized that endometrial epithelial ARID1A is critical to regulate gene expression programs necessary for early pregnancy.

In this study, we used a multi-model approach to determine the role of endometrial epithelial ARID1A in endometrial gland function and early pregnancy establishment with regard to endometriosis-related infertility. Using *Pgr*^*cre*/+^*Arid1a*^*f/f*^ mice, we found that deletion of *Arid1a* in the mouse uterus causes a gland defect starting during prepubertal development and affecting early pregnancy. ChIP analysis revealed that ARID1A directly binds at the *Foxa2* promoter at this stage. Targeting deletion of *Arid1a* to the endometrial epithelium of adult mice (*Ltf*^*iCre*/+^*Arid1a*^*f/f*^) resulted in defects of implantation, decidualization, endometrial receptivity, gland function, and critical signaling downstream of FOXA2 and LIF during early pregnancy. Furthermore, endometrial biopsy samples from women with endometriosis exhibited a decrease of FOXA2 that correlates with ARID1A levels. Decreases in both ARID1A and FOXA2 expression in the endometrium of non-human primates with induced endometriosis confirmed that this effect is endometriosis-specific.

## Materials and Methods

### Human Endometrial Tissue Samples

This study has been approved by Institutional Review Boards of Michigan State University, Greenville Health System and University of North Carolina. All methods were carried out in accordance with relevant guidelines and regulations, and written informed consent was obtained from all participants. The human endometrial samples were obtained from Michigan State University’s Center for Women’s Health Research Female Reproductive Tract Biorepository (Grand Rapids, MI), the Greenville Hospital System (Greenville, SC), and the University of North Carolina (Chapel Hill, NC). Subject selection and sample collection were performed as previously reported (20). Briefly, endometrial biopsies were obtained at the time of surgery from regularly cycling women between the ages of 18 and 45. Use of an intrauterine device (IUD) or hormonal therapies in the 3 months preceding surgery was exclusionary for this study. Histologic dating of endometrial samples was done based on the criteria of Noyes (25). For comparison of endometrium from women with endometriosis and women without endometriosis, we used eutopic endometrium derived from women with laparoscopically confirmed endometriosis and compared it to control endometrium from women laparoscopically negative for endometriosis. Control samples from 21 women were collected from the proliferative (n=7) and secretory phases (n =14). Endometriosis-affected eutopic endometrium samples from 37 women were collected from the proliferative (n=7) and secretory phases (n=30).

### Baboon Endometrium Samples

Use of the baboon endometriosis animal model was reviewed and approved by the Institutional Animal Care and Use Committees (IACUCs) of the University of Illinois at Chicago and Michigan State University. Endometriosis was induced by laparoscopically guided intraperitoneal inoculation of menstrual effluent on two consecutive cycles with a pipelle after confirmation of no pre-existing lesions as previously described (26). Eutopic endometrial tissues were collected from nine early secretory phase baboons once at pre-inoculation and again at 15-16 months post-inoculation.

### Mouse Models

All mouse procedures were approved by the Institutional Animal Care and Use Committee of Michigan State University. All mice were housed and bred in a designated animal care facility at Michigan State University under controlled humidity and temperature conditions and a 12 hour light/dark cycle at 5 mice/cage maximum. Access to water and food (Envigo 8640 rodent diet) was ad libitum. ChIP assays were conducted in C57BL/6 mice. *Arid1a* conditional knockout mice were generated in the C57BL/6 background by crossing *Pgr*^*cre*/+^(27) or *Ltf*^*iCre*/+^ (28) (The Jackson Laboratory Stock No: 026030) males with *Arid1a*^*f/f*^ (29) (generously provided by Dr. Zhong Wang, University of Michigan) females. All experiments using adult mice were performed in 8-12-week-old mice. Experiments involving *Ltf*^*iCre*/+^*Arid1a*^*f/f*^ mice were carried out using 12-week-old mice to ensure sufficient iCre activation. For breeding, one male mouse was normally placed into a cage with one female mouse. Occasionally, one male mouse was housed with two female mice to increase breeding success, and females were separated with their pups until weaning. After weaning at P21-P28, male and female littermates were housed separately until use in breeding or experiments. Mice of appropriate genotypes were randomly allocated to experimental groups, using littermates for comparisons when possible.

### Mouse Procedures and Tissue Collection

Uterine samples from specific times of pregnancy were obtained by mating control or conditional *Arid1a* knockout female mice with wildtype male mice with the morning of identification of a vaginal plug defining GD 0.5. Uteri were collected at GD 3.5, 4.5, and 5.5, and implantation sites were visualized on GD 4.5 by intravenous injection of 1% Chicago Sky Blue 6B dye (Sigma-Aldrich, St. Louis, MO) before necropsy and on GD 5.5 by gross morphology with histological confirmation. At time of dissection, isolated uterine tissue was either snap-frozen and stored at −80°C for RNA/protein extraction or fixed with 4% (vol/vol) paraformaldehyde for histological analysis. Ovaries were collected on GD 3.5 and fixed with 4% (vol/vol) paraformaldehyde for histological analysis. The serum P4 and E2 levels were analyzed by the University of Virginia Center for Research in Reproduction Ligand Assay and Analysis Core using samples taken at GD 3.5. For the fertility trial, adult female control or *Ltf*^*iCre*/+^*Arid1a*^*f/f*^ female mice were housed with wildtype male mice for 6 months, and the number of litters and pups born during that period was recorded. To quantify the number of glands, gland structures were counted based on histology in transverse tissue sections. To artificially induce decidualization, we mechanically stimulated one horn following hormonal preparation as previously described (20).

### Histology and Immunostaining

Fixed, paraffin-embedded tissues were cut at 5 μm, mounted on slides, deparaffinized, and rehydrated in a graded alcohol series. For hematoxylin and eosin staining (H&E), slides were sequentially submerged in hematoxylin, 0.25% HCL, 1% lithium carbonate, and eosin, followed by dehydration and mounting. Immunostaining was performed as previously described (20) with specific commercially available primary antibodies (Table S1). For IHC, biotinylated secondary antibodies (Vector Laboratories, Burlingame, CA) were used, followed by incubation with horseradish peroxidase (Thermo Fisher Scientific, Waltham, MA) and developing using the Vectastain Elite DAB kit (Vector Laboratories). For immunofluorescence, appropriate species-specific fluorescently tagged secondary antibodies (Invitrogen, Carlsbad, CA) were used before mounting with DAPI (Vector Laboratories) for imaging. To compare the IHC staining intensities, a semiquantitative grade (H-score) was calculated by adding the percentage of strongly stained nuclei (3x), the percentage of moderately stained nuclei (2x), and the percentage of weakly stained nuclei (1x) in a region of approximately 100 cells, giving a possible range of 0–300. The percentage of Ki67-positive cells was counted in representative fields of approximately 150 epithelial cells and 150 stromal cells.

### Whole Mount Immunofluorescence and Imaging

Three-dimensional imaging of mouse uteri was performed as previously described (30). Briefly, samples were fixed with a 4:1 ratio of Methanol:DMSO and stained using whole-mount immunofluorescence. Primary antibodies (Table S1) for mouse CDH1 (E-cadherin) and FOXA2 were utilized to identify total epithelium and glandular epithelium, respectively, followed by fluorescently conjugated Alexa Fluor IgG (Invitrogen) secondary antibodies. Embryos were identified using a combination of Hoechst and CDH1 (E-cadherin). Uteri were imaged using a Leica SP8 TCS confocal microscope with white-light laser, using a 10× air objective with *z* stacks that were 7 μm apart. Full uterine horns were imaged using 18×2 tile scans and tiles were merged using the mosaic merge function of the Leica software. LIF files (Leica software) were analyzed using Imaris v9.1 (Bitplane, Zürich, Switzerland). Using the channel arithmetic function, the glandular FOXA2+ signal was removed from the E-CAD+ signal to create lumen-only signal. Surfaces were created in surpass 3D mode for the lumen-only signal and the FOXA2+ glandular signal.

### RNA Isolation and RT-qPCR

As previously described (20), RNA was extracted from the uterine tissues using the RNeasy total RNA isolation kit (Qiagen, Valencia, CA). mRNA expression levels were measured by real-time PCR TaqMan or SYBR green analysis using an Applied Biosystems StepOnePlus system according to the manufacturer's instructions (Applied Biosystems, Foster City, CA) using pre-validated primers (Table S2), probes (Table S3), and either PowerUp SYBR Green Master Mix or TaqMan Gene Expression Master Mix (Applied Biosystems). Template cDNA was produced from 3 μg of total RNA using random hexamers and MMLV Reverse transcriptase (Invitrogen). The mRNA quantities were normalized against Rpl7 for SYBR green primers or Gapdh for TaqMan probes.

### Chromatin Immunoprecipitation

ChIP assays were conducted by Thermo Scientific (Pittsburgh, PA, USA) using uteri of C57BL/6 mice at day 0.5 and 3.5 of gestation (GD 0.5 and 3.5). ChIP assays were performed as previously described (20). Briefly, for each ChIP reaction, 100 μg of chromatin was immunoprecipitated by 4 μg of antibodies against ARID1A (Table S1). Eluted DNA was amplified with specific primers (Table S2) using SYBR Green Master (Roche, Basel, Switzerland), and the resulting signals were normalized to input activity.

### Comparative Transcriptomic Analysis

Comparisons of GD 3.5 *Pgr*^*cre*/+^*Arid1a*^*f/f*^ to GD 3.5 *Pgr*^*cre*/+^*Foxa2*^*f/f*^ uterine dysregulated genes were performed by comparing differentially expressed genes determined in our previously reported, publicly available *Pgr*^*cre*/+^*Arid1a*^*f/f*^ transcriptomics analysis (GSE72200) (20) with differentially expressed genes determined by previously reported transcriptomics analysis of *Pgr*^*cre*/+^*Foxa2*^*f/f*^ mice publicly available from the GEO database (GSE48339) (31). Before calculating overlap, duplicate genes were removed based on GeneBank ID or gene symbol.

### Statistical Analysis

To assess statistical significance of parametric data, we used student’s t-test for comparison of two groups or one-way ANOVA followed by Tukey’s post-hoc test for multiple comparisons. For non-parametric data, we used the Mann-Whitney U test for comparison of two groups or Kruskal-Wallis rank test followed by Dunn’s test for multiple comparisons. Pearson’s Correlation Coefficient was used to assess correlation. All statistical tests were two-tailed when applicable, and a value of p<0.05 was considered statistically significant. Statistical analyses were performed using either the Instat or Prism package from GraphPad (San Diego, CA). Statistical test results (p-values) are presented with the results in the text and symbolically in the figures, with explanations in figure legends. The value of n for each experiment, representing number of animals unless noted as number of implantation sites, is reported in the appropriate figure legend.

### Data and Code Availability

The data that support the findings of this study are available from the article and Supplementary Information files.

## Results

### Uterine ARID1A is Critical for Endometrial Gland Development and Function and Binds the *Foxa2* Promoter in Pregnant Mice

Because of the importance of FOXA2 and endometrial glands in the implantation process (10, 13, 32), we analyzed gland formation and function in uterine-specific *Arid1a* knockout mice (*Pgr*^*cre*/+^*Arid1a*^*f/f*^) (20). *Pgr*^*cre*/+^*Arid1a*^*f/f*^ mice had significantly fewer endometrial glands compared to controls at gestation day (GD) 3.5, which marks the pre-implantation stage (3) (−2.20 fold, p=0.0165; Fig. 1*A*). Immunohistochemical (IHC) analysis revealed a lack of FOXA2 in *Pgr*^*cre*/+^*Arid1a*^*f/f*^ glands (Fig. 1*B*). Due to the limitations of analyzing thin tissue sections, we utilized a whole-mount immunofluorescence approach combined with confocal imaging and quantitative image analysis to visualize the uterine structure in three dimensions (30). Three-dimensional imaging during early pregnancy revealed that in contrast to the abundance of uniformly oriented FOXA2-positive glands in control mice, *Pgr*^*cre*/+^*Arid1a*^*f/f*^ mice exhibited very few randomly scattered FOXA2-positive glands (Fig. 1*C*). In order to more clearly understand the molecular dysregulation, we utilized our previously published transcriptomic data from GD 3.5 *Pgr*^*cre*/+^*Arid1a*^*f/f*^ mouse uteri (20) to compare the dysregulated genes to those dysregulated in *Pgr*^*cre*/+^*Foxa2*^*f/f*^ uteri at GD 3.5 (31). Out of a total of 2,075 genes differentially expressed due to *Arid1a* loss (2,556 probes >1.5 fold change, duplicate genes removed), 316 (15.23%) were also differentially expressed in *Pgr*^*cre*/+^*Foxa2*^*f/f*^ mice (out of 915 probes >1.5 fold change, duplicate genes removed; Fig. 1*D*, Dataset S1). Due to the importance of LIF in implantation (33) and its diminished expression in the GD 3.5 *Pgr*^*cre*/+^*Foxa2*^*f/f*^ mouse uterus (10, 13), we analyzed *Lif* expression in the GD 3.5 *Pgr*^*cre*/+^*Arid1a*^*f/f*^ mouse uterus with RT-qPCR and found it to be significantly reduced (−8.44 fold, p=0.0357; Fig. 1*E*). We also confirmed that *Foxa2* mRNA transcripts were decreased in the *Pgr*^*cre*/+^*Arid1a*^*f/f*^ mouse uterus along with *Spink3* and *Cxcl15*, previously recognized gland-specific genes (10, 31, 32) (−8.83 fold, p=0.0159; −71.86 fold, p=0.0159; −5.66 fold, p=0.0002; respectively; Fig. 1*E*). Together, these findings reveal a major defect of endometrial gland structure and function in *Pgr*^*cre*/+^*Arid1a*^*f/f*^ mice during early pregnancy.

**Figure 1.**
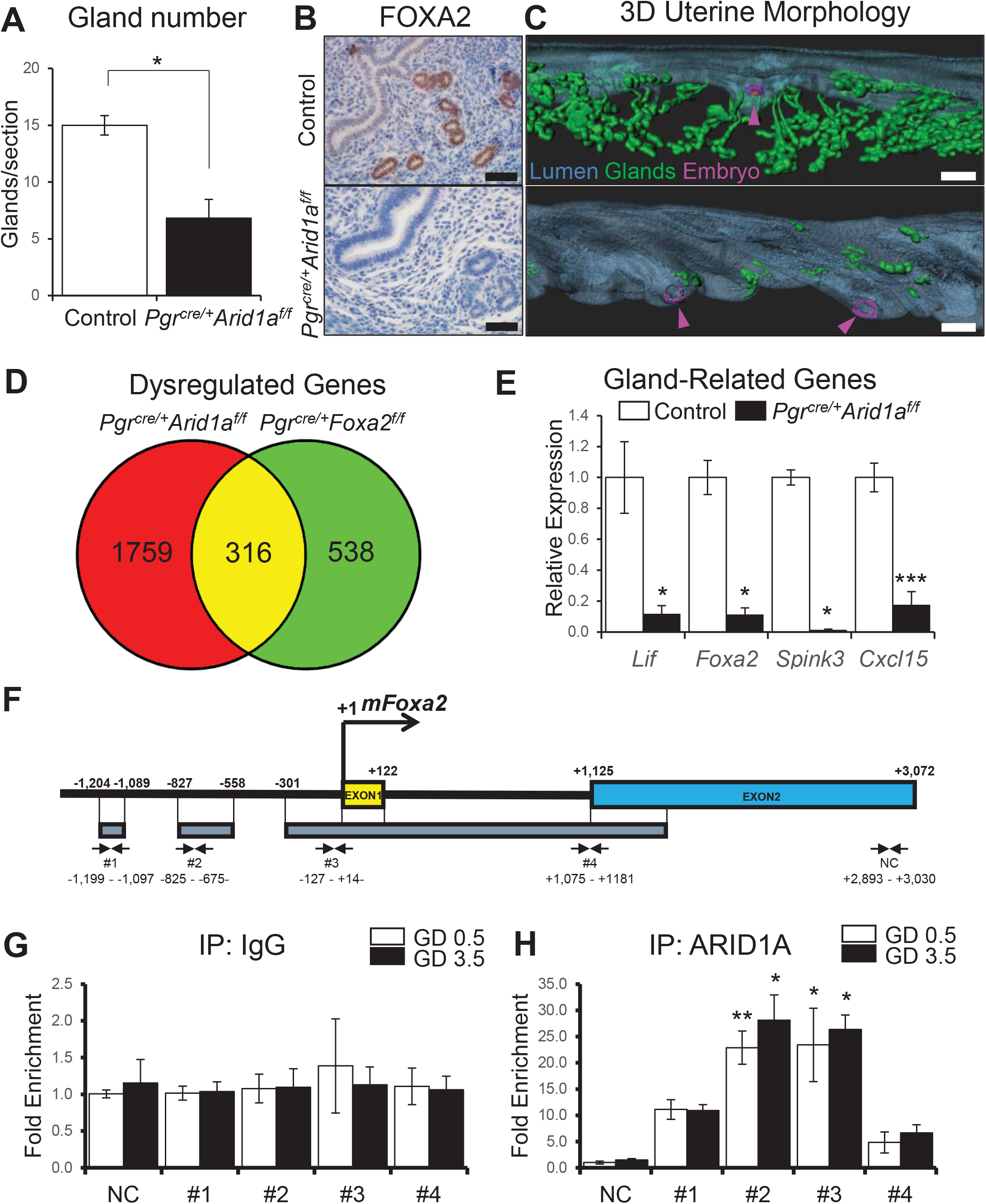
Uterine ARID1A is Critical for Endometrial Gland Development and Function and Binds the *Foxa2* Promoter in Pregnant Mice. *(A)* Endometrial gland counts in control and *Pgr*^*cre*/+^*Arid1a*^*f/f*^ mice at GD 3.5. The graph represents the mean ± SEM of the number of glands per uterine tissue section (n=6; *, p<0.05). *(B)* Representative images of FOXA2 IHC in control and *Pgr*^*cre*/+^*Arid1a*^*f/f*^ mouse uterine sections at GD 3.5 (n=4). Scale bars: 50 μm. (*C)* Three-dimensional uterine morphology of control and *Pgr*^*cre*/+^*Arid1a*^*f/f*^ uterine horns during early pregnancy based on whole-mount immunofluorescence for E-cadherin and FOXA2, where the 3D luminal structure (blue) is constructed by subtracting the FOXA2 (green) from the E-cadherin signal (n=4). Arrowheads indicate embryos within the uterine horns. Scale bars: 200 μm. *(D)* Overlapping genes dysregulated at GD 3.5 in the uterus by deletion of *Arid1a* or *Foxa2*. *(E)* Relative expression of endometrial gland-related gene mRNA normalized to *Gapdh* (*Lif*, *Foxa2*) or *Rpl7* (*Spink 3*, *Cxcl15*) in whole uterine tissue preparations at GD 3.5. The graphs represent the mean ± SEM (Control, n=35; *Pgr*^*cre*/+^*Arid1a*^*f/f*^, n=5); *, p<0.05; ***, p<0.001). *(F)* The schematic shows putative ARID1A binding sites near the *Foxa2* gene (#1, 2, 3, 4). *(G)* Fold enrichment based on RT-qPCR targeting putative ARID1A binding sites in GD 0.5 and GD 3.5 mouse uteri after ChIP using IgG control. The graph shows the mean ± SEM (n=5). *(H)* Fold enrichment based on RT-qPCR targeting putative ARID1A binding sites in GD 0.5 and GD 3.5 mouse uteri after ChIP using anti-ARID1A antibody. The graph shows the mean ± SEM (n=5; *, p<0.05; **, p<0.01).

Because deletion of *Foxa2* during early postnatal development causes drastic loss of gland formation (13, 31), we analyzed the 3-4-week-old *Pgr*^*cre*/+^*Arid1a*^*f/f*^ mouse uterus to determine if the gland defect found during early pregnancy was also present before maturity. Indeed, 4-week-old *Pgr*^*cre*/+^*Arid1a*^*f/f*^ mice exhibited a significantly decreased endometrial gland number and loss of FOXA2 expression, and 3-week-old *Pgr*^*cre*/+^*Arid1a*^*f/f*^ mice showed a lack of FOXA2-positive gland elongation compared to controls based on 3D image reconstruction (Fig. S1*A-C*). These findings indicate that the structural and functional gland defect in *Pgr*^*cre*/+^*Arid1a*^*f/f*^ mice is not specific to early pregnancy but starts during prepubertal development.

Analysis of publicly available ARID1A ChIP-seq data from HepG2 cells (34) identified putative binding sites for ARID1A near the *Foxa2* gene (Fig. 1*F*). To determine whether ARID1A binds these sites in the mouse uterus during early pregnancy, we performed ChIP-qPCR on whole uterine tissue lysates from wildtype mice at GD 0.5 and GD 3.5 with primers designed to target the putative binding regions. As expected, immunoprecipitation (IP) with nonspecific IgG showed no region of significant enrichment compared to the negative control; however, IP with an ARID1A antibody followed by qPCR revealed significant enrichment of putative binding sites #2 (GD 0.5, 22.95 fold, p<0.01; GD 3.5, 18.73 fold, p<0.05) and #3 (GD 0.5, 23.47 fold, p<0.05; GD 3.5, 17.53 fold, p<0.05) over the negative control region (Fig. 1*G-H*). This finding indicates that ARID1A directly binds the *Foxa2* promoter region in vivo.

### Endometrial Epithelial-Specific *Arid1a* Loss Causes Severe Sub-Fertility and Compromises Gland Function in Mice

To further dissect the relationship between ARID1A and FOXA2 in the endometrium during early pregnancy, we conditionally ablated *Arid1a* in the adult mouse endometrial epithelium by crossing *Ltf*^*iCre*/+^ (28) and *Arid1a*^*f/f*^ (29) mice. When crossed with *Arid1a*^*f/f*^ mice, this model causes *Arid1a* deletion in the luminal and glandular epithelium, while retaining *Arid1a* expression in the stroma in contrast to *Pgr*^*cre*/+^*Arid1a*^*f/f*^ mice which delete *Arid1a* in both epithelial and stromal compartments (Fig. 2*A-B*). Additionally, this approach circumvents any developmental defects by restricting iCre expression to adulthood. To assess overall fecundity, *Ltf*^*iCre*/+^*Arid1a*^*f/f*^ and control females were housed with wildtype male mice in a six month fertility trial. Both groups of mice engaged in normal mating activity resulting in the observation of copulatory plugs, and control mice had expected numbers of litters and pups/litter. However, *Ltf*^*iCre*/+^*Arid1a*^*f/f*^ females were found to be severely sub-fertile, and the only three pups found were dead upon discovery (n=6; Fig. S2*A*). To determine if an ovarian defect was responsible for the severe sub-fertility of *Ltf*^*iCre*/+^*Arid1a*^*f/f*^ females, we examined ovarian histology, finding no anatomical abnormalities and normal development of corpora lutea (Fig. S2*B*). Furthermore, analysis of serum ovarian steroid hormone levels at GD 3.5 revealed no differences between *Ltf*^*iCre*/+^*Arid1a*^*f/f*^ mice and controls in total E2 or P4 (Fig. S2*C*). These analyses indicate that the severe sub-fertility phenotype of *Ltf*^*iCre*/+^*Arid1a*^*f/f*^ mice is not due to a defect of ovarian function.

**Figure 2.**
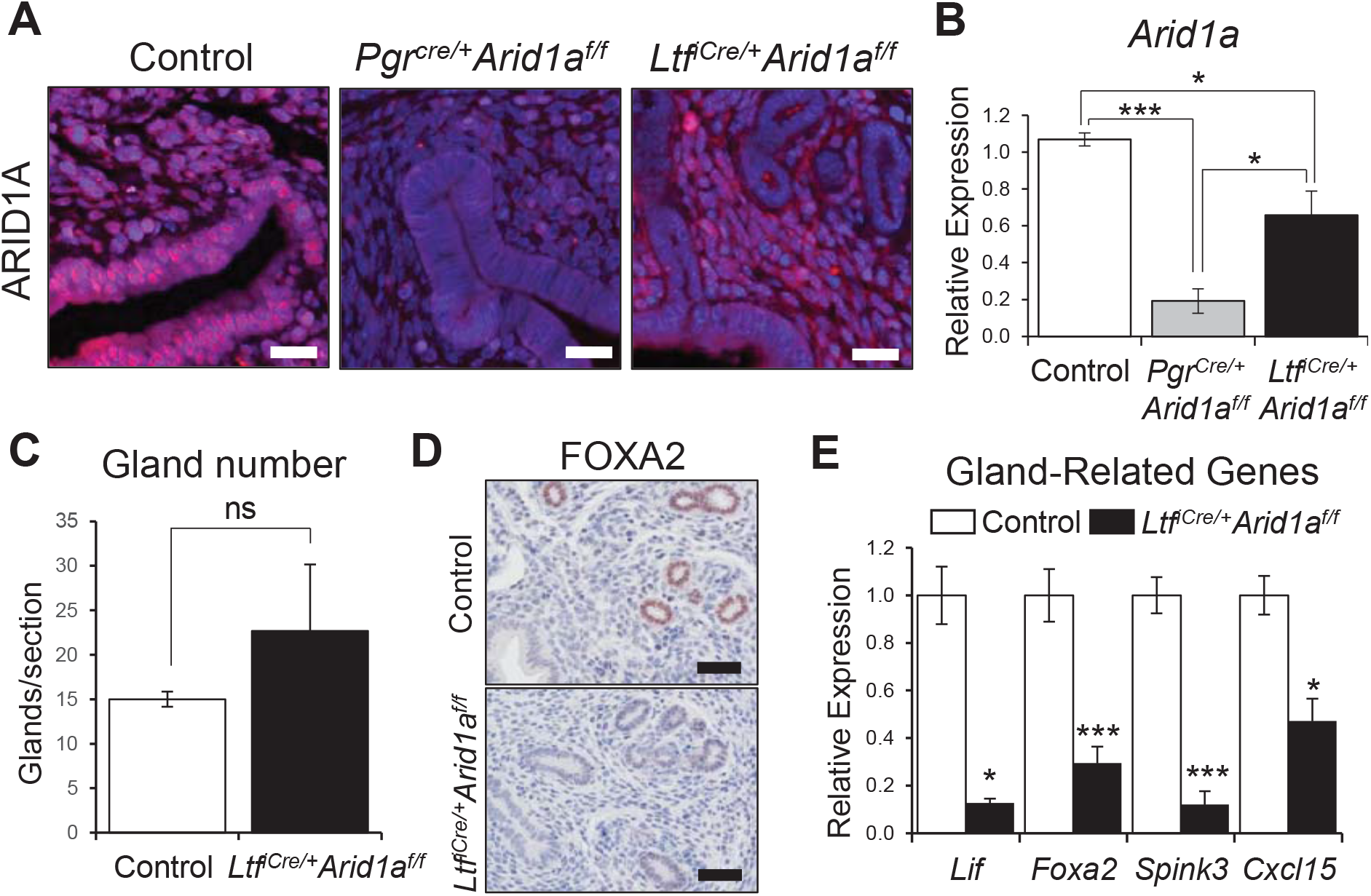
Endometrial Epithelial-Specific *Arid1a* Loss Compromises Gland Function. *(A)* Representative images show immunofluorescence staining of ARID1A (Texas Red) and DAPI (Blue) demonstrating strong ARID1A expression in the endometrial epithelium and stroma of control mice, loss of ARID1A expression in both epithelium and stroma of *Pgr*^*cre*/+^*Arid1a*^*f/f*^ mice, and loss of ARID1A in the epithelium but not stroma of *Ltf*^*iCre*/+^*Arid1a*^*f/f*^ mice at GD 3.5 (n=3/genotype). Scale bars: 25 μm. *(B)* Relative expression of *Arid1a* mRNA normalized to *Rpl7* in RT-qPCR using whole uterine tissue preparations. The graph displays the mean ± SEM (n=3/genotype; *, p<0.05; ***, p<0.001). *(C)* Endometrial gland counts in control and *Ltf*^*iCre*/+^*Arid1a*^*f/f*^ mice at GD 3.5. The graph represents the mean ± SEM of the number of glands per uterine tissue section (n=6; ns, p>0.05). *(D)* Representative images of FOXA2 IHC in control and *Ltf*^*iCre*/+^*Arid1a*^*f/f*^ mouse uterine sections (n=4). Scale bars: 50 μm. *(E)* Relative expression of endometrial gland-related gene mRNA normalized to *Rpl7* in whole uterine tissue preparations determined with RT-qPCR. The graphs represent the mean ± SEM (Control, n=45; *Ltf*^*iCre*/+^*Arid1a*^*f/f*^, n=5; *, p<0.05; ***, p<0.001).

To determine if endometrial epithelial *Arid1a* loss in adult mice compromises gland structure or function, we examined *Ltf*^*iCre*/+^*Arid1a*^*f/f*^ endometrial glands at GD 3.5. Gland counts from transverse uterine tissue sections revealed no difference in gland number between *Ltf*^*iCre*/+^*Arid1a*^*f/f*^ mice and controls (Fig. 2*C*). Though the quantity of glands was unchanged, *Ltf*^*iCre*/+^*Arid1a*^*f/f*^ mice exhibited a defect of gland function at GD 3.5 indicated by decreased FOXA2 expression and significant decreases in uterine *Lif* (−7.89 fold, p=0.0159), *Foxa2* (−3.38 fold, p<0.0001), *Spink3* (−8.26 fold, p<0.0001), and *Cxcl15* (−3.12 fold, p=0.0159) mRNA levels (Fig. 2*D-E*).

### *Ltf*^*iCre*/+^*Arid1a*^*f/f*^ Mice Exhibit Implantation and Decidualization Defects

On the morning of GD 4.5, implantation sites were visible in control mouse uteri but not in *Ltf*^*iCre*/+^*Arid1a*^*f/f*^ uteri (Fig. 3*A*). Hematoxylin and eosin (H&E) staining of transverse tissue sections revealed that though luminal closure and embryo apposition had occurred in *Ltf*^*iCre*/+^*Arid1a*^*f/f*^ mice, the decidualization response of stromal cells surrounding the embryo did not occur as in controls (Fig. 3*B*). COX2, a marker of decidualization (10), was present in stromal cells surrounding the embryo in controls but was limited to the epithelium in *Ltf*^*iCre*/+^*Arid1a*^*f/f*^ mice, providing molecular evidence of early decidualization response failure (Fig. 3*C*).

**Figure 3.**
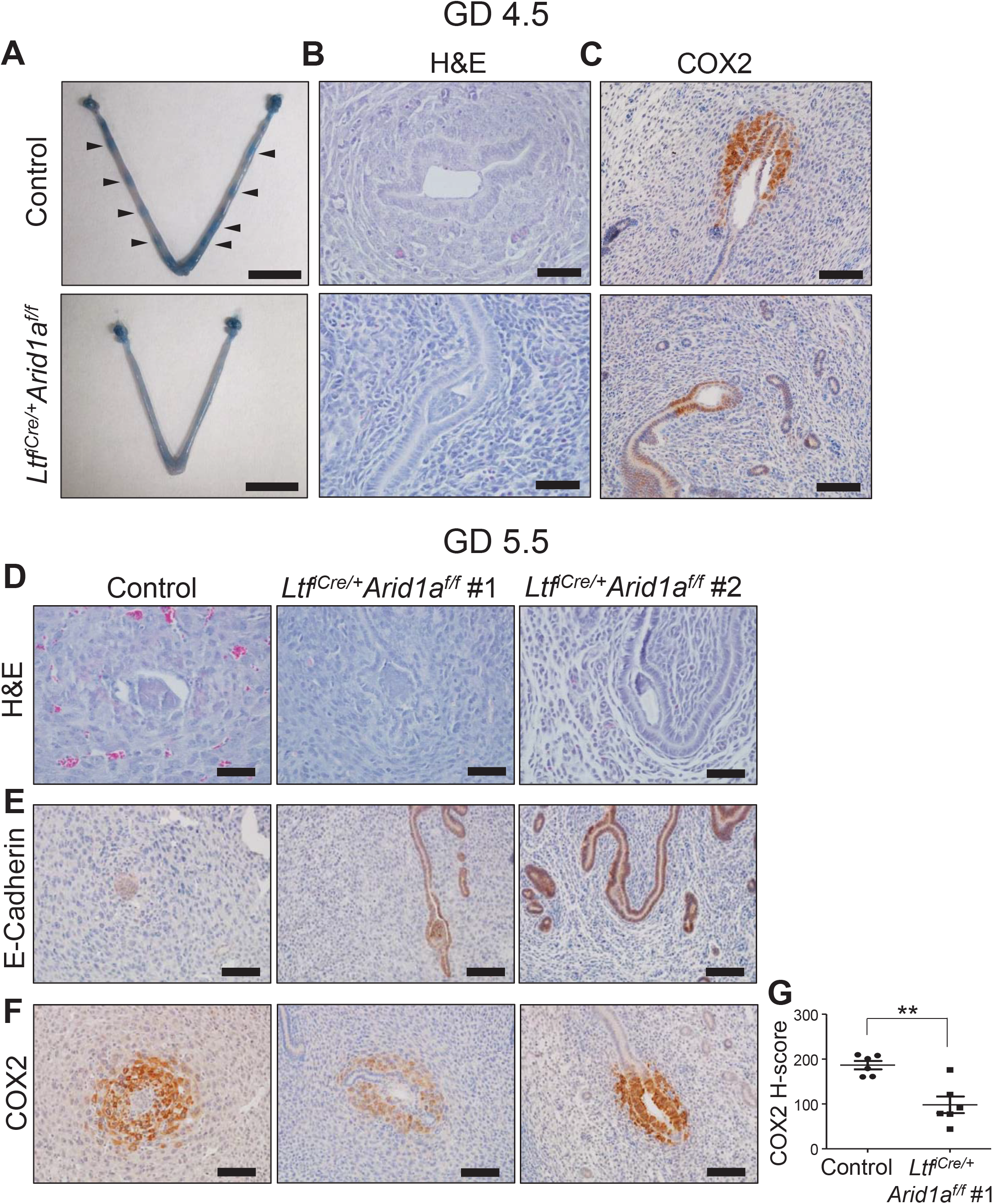
*Ltf*^*iCre*/+^*Arid1a*^*f/f*^ Mice Exhibit an Implantation Defect. *(A)* Implantation sites were grossly visible in control but not *Ltf*^*iCre*/+^*Arid1a*^*f/f*^ uteri at GD 4.5 after intravenous injection of Chicago Sky Blue 6B dye. Arrowheads indicate clear implantation sites (IS; n=3). Scale bars: 1 cm. *(B)* Representative images of hematoxylin and eosin (H&E) staining showing decidualization of stromal cells around an implanting embryo in control but not *Ltf*^*iCre*/+^*Arid1a*^*f/f*^ uterine sections at GD 4.5 (n=11 IS). Scale bars: 50 μm. *(C)* Representative images of COX2 IHC in control and *Ltf*^*iCre*/+^*Arid1a*^*f/f*^ mouse uteri at GD 4.5 (n=6 IS). Scale bars: 100 μm. *(D)* Representative images of hematoxylin and eosin (H&E) staining in control and *Ltf*^*iCre*/+^*Arid1a*^*f/f*^ uterine sections at GD 5.5 (n=5). In contrast to the control, the *Ltf*^*iCre*/+^*Arid1a*^*f/f*^ mouse luminal epithelium remained intact surrounding the embryos. Most *Ltf*^*iCre*/+^*Arid1a*^*f/f*^ implantation sites (IS; 18/21, 86%) exhibited some decidualizing stromal cells based on morphology (*Ltf*^*iCre*/+^*Arid1a*^*f/f*^ #1), though not to the same degree as controls. Some *Ltf*^*iCre*/+^*Arid1a*^*f/f*^ IS (3/21, 14%), however, underwent little to no decidualization (*Ltf*^*iCre*/+^*Arid1a*^*f/f*^ #2). Scale bars: 50 μm. *(E)* Representative images of E-cadherin IHC in control and *Ltf*^*iCre*/+^*Arid1a*^*f/f*^ mouse uteri at GD 5.5 (control and *Ltf*^*iCre*/+^*Arid1a*^*f/f*^ #1, n=6 IS; *Ltf*^*iCre*/+^*Arid1a*^*f/f*^ #2, n=3 IS). Scale bars: 100 μm. *(F)* Representative images of COX2 IHC in control and *Ltf*^*iCre*/+^*Arid1a*^*f/f*^ mouse uteri at GD 5.5 (control and *Ltf*^*iCre*/+^*Arid1a*^*f/f*^ #1, n=6 IS; *Ltf*^*iCre*/+^*Arid1a*^*f/f*^ #2, n=3 IS). Scale bars: 100 μm. *(G)* Semi-quantitative H-score of COX2 staining strength in uterine sections of control versus *Ltf*^*iCre*/+^*Arid1a*^*f/f*^ #1 phenotype group mice at GD 5.5. The graph represents the mean ± SEM (n=6 IS; **, p<0.01).

At post-implantation (GD 5.5), H&E analysis and IHC for E-cadherin, an epithelial tight junction protein, showed that the luminal epithelium remained intact around the embryo in all *Ltf*^*iCre*/+^*Arid1a*^*f/f*^ mice analyzed, whereas no luminal epithelium remained in controls due to successful embryo implantation (Fig. 3*D-E*). H&E and COX2 IHC revealed two subsets of decidual cell phenotypes in *Ltf*^*iCre*/+^*Arid1a*^*f/f*^ mice at GD 5.5. In 85.71% (18/21) of implantation sites observed (*Ltf*^*iCre*/+^*Arid1a*^*f/f*^ #1), some decidualization occurred, but not to the degree of controls as evidenced by less apparent decidual cell morphology and a significant reduction in COX2 expression as indicated by H-score (−1.90 fold, p=0.0017; Fig. 3*D-F*, center panel; *G*). In the remaining 14.29% of implantation sites (*Ltf*^*iCre*/+^*Arid1a*^*f/f*^ #2), little to no decidualization was apparent in the stroma (Fig. 3*D-F*, right panel). In spite of defective implantation, embryos in *Ltf*^*iCre*/+^*Arid1a*^*f/f*^ uteri were firmly attached and could not be flushed (n=3).

To confirm the decidualization defect of *Ltf*^*iCre*/+^*Arid1a*^*f/f*^ mice, we performed hormonally and physically stimulated artificial decidualization induction to mimic an invading embryo (20). Though the stimulated horn of control mice underwent a robust decidualization reaction by decidualization day 5, very little decidualization occurred in the stimulated horn of *Ltf*^*iCre*/+^*Arid1a*^*f/f*^ mice as evidenced by gross morphology, significantly decreased uterine horn weight ratio (−4.80 fold, p=0.0022), and cell morphology (Fig. 4). This result clearly confirmed that the lack of a proper decidualization response in *Ltf*^*iCre*/+^*Arid1a*^*f/f*^ endometrial stromal cells was not merely a result of a lack of stimulation by the embryo. Together, these data display an implantation defect in *Ltf*^*iCre*/+^*Arid1a*^*f/f*^ mice resulting primarily from a failure of decidualization response.

**Figure 4.**
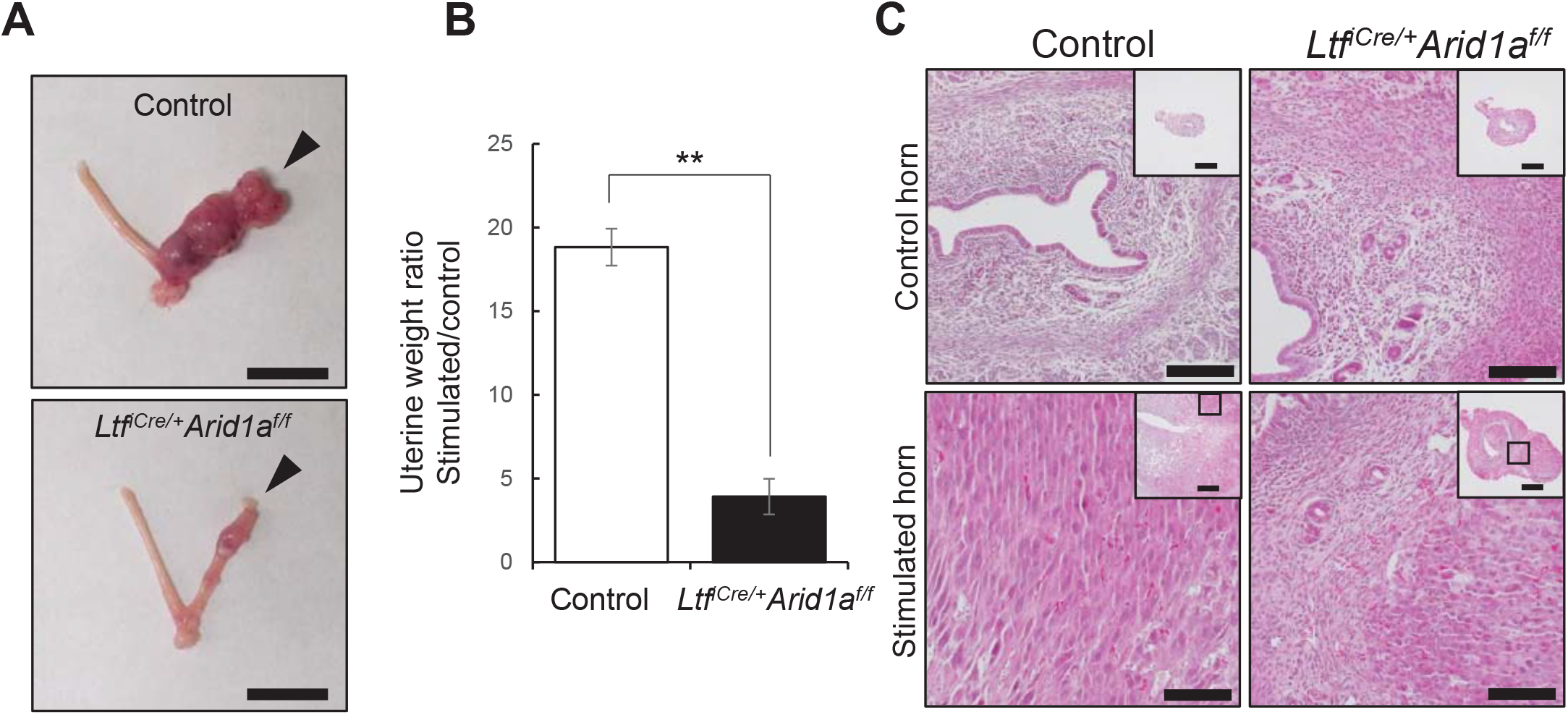
*Ltf*^*iCre*/+^*Arid1a*^*f/f*^ Mice Exhibit a Decidualization Defect. *(A)* Gross morphology of decidualization day 5 control and *Ltf*^*iCre*/+^*Arid1a*^*f/f*^ uteri, arrowheads indicating the stimulated uterine horn (n=6). Scale bars: 1 cm. *(B)* Stimulated/control horn weight ratio in control and *Ltf*^*iCre*/+^*Arid1a*^*f/f*^ mice. The graph represents the mean ± SEM (n=6; **, p<0.01). *(C)* Representative images of hematoxylin and eosin staining show the decidual cell morphology in the stimulated horn of controls but defective decidualization in *Ltf*^*iCre*/+^*Arid1a*^*f/f*^ mice (n=6). Scale bars: main images 100 μm, insets 500 μm.

### *Ltf*^*iCre*/+^*Arid1a*^*f/f*^ Mice Exhibit a Non-Receptive Endometrium

To assess the receptivity of the luminal epithelium in *Ltf*^*iCre*/+^*Arid1a*^*f/f*^ mice, we analyzed cell proliferation at pre-implantation (GD 3.5) using IHC for Ki67 and found that epithelial cell proliferation was significantly increased (16.44 fold, p<0.001) and stromal cell proliferation was significantly decreased (−4.71 fold, p=0.001) in *Ltf*^*iCre*/+^*Arid1a*^*f/f*^ uteri compared to controls (Fig. 5). A highly proliferative epithelium at this stage indicates a non-receptivity to embryo implantation (20), while an under-proliferative stroma could indicate a failure to properly prepare for decidualization (1, 3).

**Figure 5.**
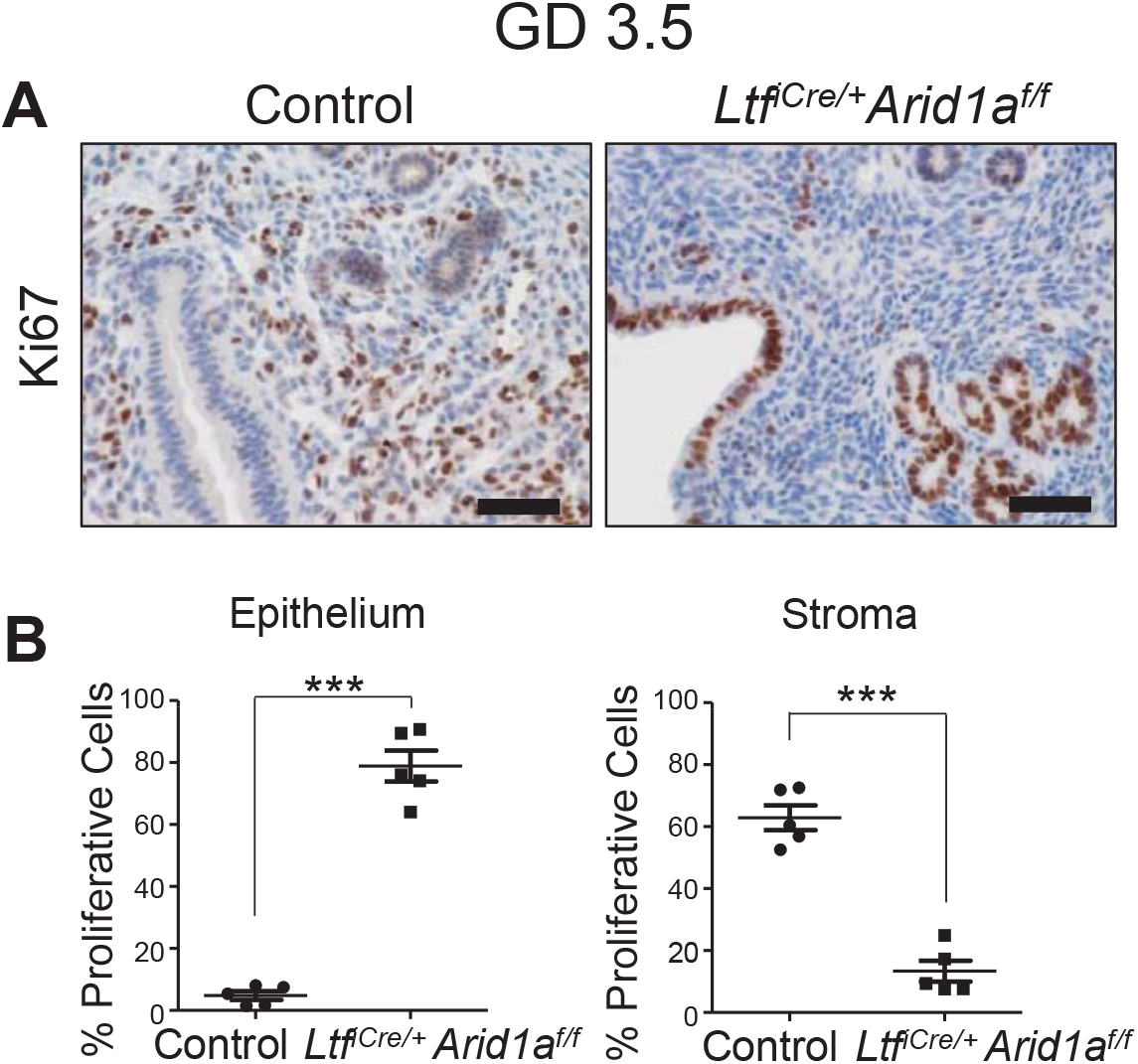
*Ltf*^*iCre*/+^*Arid1a*^*f/f*^ Mice Exhibit a Non-Receptive Endometrium. *(A)* Representative images of Ki67 IHC in control and *Ltf*^*iCre*/+^*Arid1a*^*f/f*^ mouse uteri (n=3). Scale bars: 50 μm. *(B)* Percentage of proliferative cells in epithelial and stromal compartments of mouse endometrial tissue sections. The graphs represent the mean percentage of proliferative cells ± SEM (n=3, 5 tissue regions; ***, p<0.001).

A loss of PGR as previously shown in *Pgr*^*cre*/+^*Arid1a*^*f/f*^ mice (20) could explain the finding of a highly proliferative epithelium. However, the *Ltf*^*iCre*/+^*Arid1a*^*f/f*^ epithelium largely retained PGR expression at GD 3.5, and the PGR signal strength was not significantly different from controls based on H-score in the epithelium or the stroma, although there were patches of epithelial cells negative for PGR (Fig. S3*A-B*). Western blotting confirmed that total levels of the PGR isoforms PR-A and PR-B were not changed in *Ltf*^*iCre*/+^*Arid1a*^*f/f*^ uteri (Fig. S3*C-D*). Correspondingly, IHC revealed no changes in total ESR1 or pESR1 levels at this stage in *Ltf*^*iCre*/+^*Arid1a*^*f/f*^ uterine sections (Fig. S3*E-F*). RT-qPCR analysis of P4 and E2 target gene expression in GD 3.5 *Ltf*^*iCre*/+^*Arid1a*^*f/f*^ uteri revealed that though the majority of genes tested were not different than controls to the level of statistical significance, the P4 targets *Areg* and *Lrp2* (35) were significantly decreased, and the E2 targets *Clca3*, *Ltf*, and *C3* were significantly increased (Fig. S3*G-H*). Taken together, these results indicate that PGR loss in patches of epithelial cells likely contributes to increased E2-induced epithelial proliferation in *Ltf*^*iCre*/+^*Arid1a*^*f/f*^ mice, though not to the degree previously noted in *Pgr*^*cre*/+^*Arid1a*^*f/f*^ mice (20).

FOXO1 is a transcription factor with known roles in regulating epithelial integrity (36) and decidualization (37) during implantation, and its expression is reciprocal to PGR (36). We profiled FOXO1 and PGR expression in *Ltf*^*iCre*/+^*Arid1a*^*f/f*^ mice during early pregnancy using IHC. At GD 3.5, we found that in patches of the luminal epithelium negative for PGR expression, nuclear FOXO1 expression was increased (Fig. S4*A*). Observation of the epithelium around implantation sites (IS) at GD 4.5 and at GD 5.5 inter-implantation sites (I-IS) revealed no marked changes in FOXO1 or PGR expression between *Ltf*^*iCre*/+^*Arid1a*^*f/f*^ mice and controls, implying that the non-receptive endometrium of *Ltf*^*iCre*/+^*Arid1a*^*f/f*^ mice is not due to dysregulation of FOXO1 (Fig. S4*B-C*).

### The pSTAT3-EGR1 Pathway is Dysregulated during Early Pregnancy in *Ltf*^*iCre*/+^*Arid1a*^*f/f*^ Mice

Since *Ltf*^*iCre*/+^*Arid1a*^*f/f*^ mice exhibited decreased *Lif* expression at the pre-implantation stage, we examined the downstream effects of the decrease in *Lif* as another potential explanation for the defects in implantation, decidualization, and endometrial receptivity. For successful implantation to take place, signal transducer and activator of transcription 3 (STAT3) must be activated by phosphorylation (pSTAT3) downstream of LIF after the GD 3.5 E2 surge, whereupon it induces early growth response 1 (EGR1) expression in the stroma (38–40). At GD 3.5, pSTAT3 is normally expressed robustly in endometrial epithelial cells (38), but IHC analysis revealed that *Ltf*^*iCre*/+^*Arid1a*^*f/f*^ mice exhibit significantly decreased pSTAT3 (−8.96 fold, p=0.0079; Fig. 6*A-B*). The further dysregulation of this pathway was evident in that EGR1 expression was significantly reduced in the GD 3.5 *Ltf*^*iCre*/+^*Arid1a*^*f/f*^ endometrial stroma based on IHC H-score (−2.09 fold, p=0.0242; Fig. 6*C-D*).

**Figure 6.**
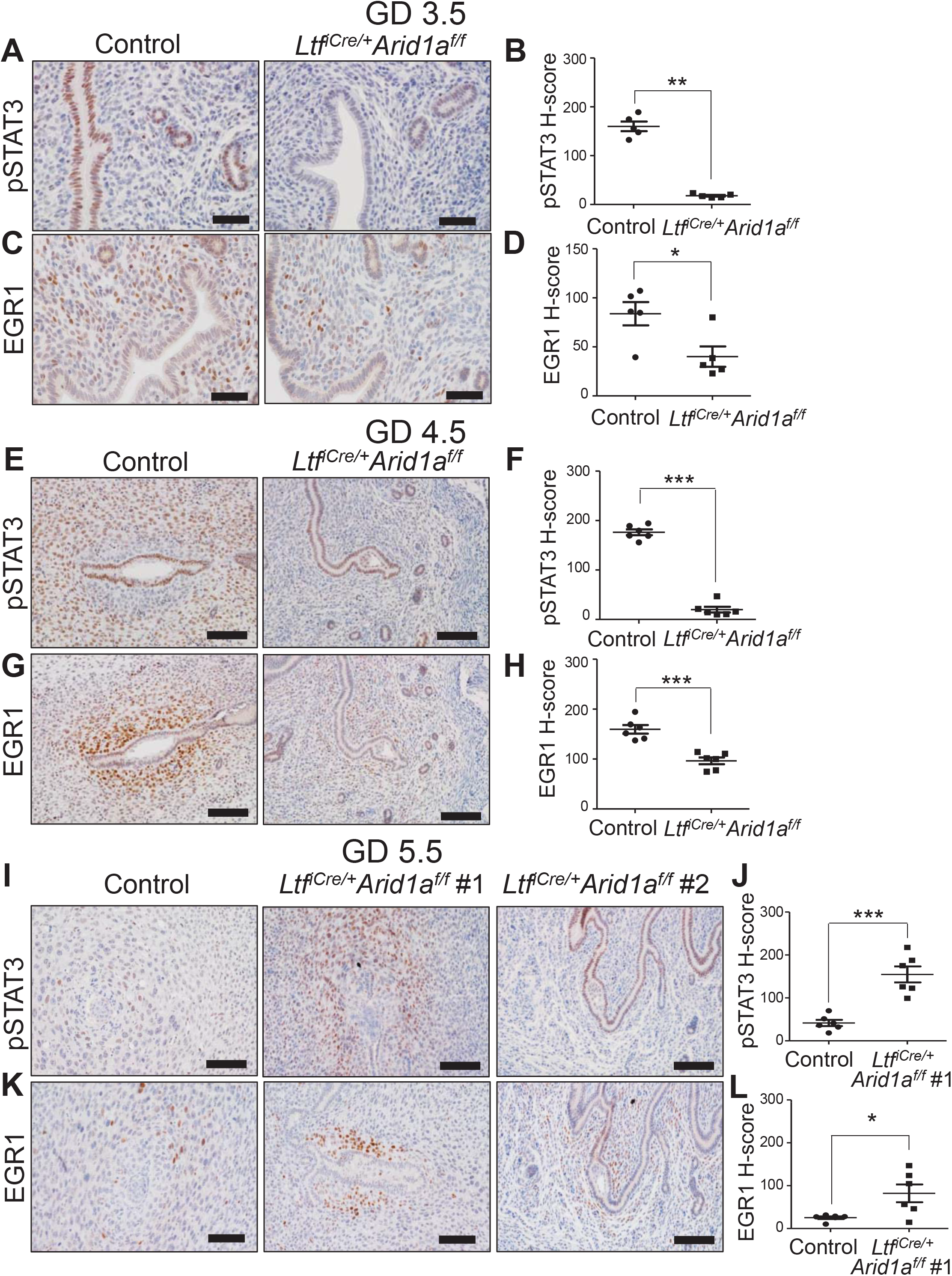
The pSTAT3-EGR1 Pathway is Dysregulated during Early Pregnancy in *Ltf*^*iCre*/+^*Arid1a*^*f/f*^ Mice. *(A)* Representative images of pSTAT3 IHC in control and *Ltf*^*iCre*/+^*Arid1a*^*f/f*^ mouse uterine tissue sections at GD 3.5 (n=3). Scale bars: 50 μm. *(B)* Semi-quantitative H-score of epithelial pSTAT3 staining strength at GD 3.5. The graph represents the mean ± SEM (n=3, 5 tissue regions; **, p<0.01). *(C)* Representative images of EGR1 IHC in control and *Ltf*^*iCre*/+^*Arid1a*^*f/f*^ mouse uterine tissue sections at GD 3.5 (n=3). Scale bars: 50 μm. *(D)* Semi-quantitative H-score of stromal EGR1 staining strength. The graph represents the mean ± SEM (n=3, 5 tissue regions; *, p<0.05). *(E)* Representative images of pSTAT3 IHC in control and *Ltf*^*iCre*/+^*Arid1a*^*f/f*^ mouse uterine tissue sections at GD 4.5 (n=6 IS). Scale bars: 100 μm. *(F)* Semi-quantitative H-score of stromal pSTAT3 staining strength. The graph shows the mean ± SEM (n= 6 IS; ***, p<0.001). *(G)* Representative images of EGR1 IHC in control and *Ltf*^*iCre*/+^*Arid1a*^*f/f*^ mouse uterine tissue sections at GD 4.5 (n=6 IS). Scale bars: 100 μm. *(H)* Semi-quantitative H-score of stromal EGR1 staining strength. The graph shows the mean ± SEM (n=6 IS; ***, p<0.001). *(I)* Representative images of pSTAT3 IHC in control and *Ltf*^*iCre*/+^*Arid1a*^*f/f*^ mouse uterine tissue sections at GD 5.5 (control and *Ltf*^*iCre*/+^*Arid1a*^*f/f*^ #1, n=6 IS; *Ltf*^*iCre*/+^*Arid1a*^*f/f*^ #2, n= 3 IS). Scale bars: 100 μm. *(J)* Semi-quantitative H-score of stromal pSTAT3 staining strength in control versus *Ltf*^*iCre*/+^*Arid1a*^*f/f*^ #1. The graph shows the mean ± SEM (n=6 IS; ***, p<0.001). (*K*) Representative images of EGR1 IHC in control and *Ltf*^*iCre*/+^*Arid1a*^*f/f*^ mouse uterine tissue sections at GD 5.5 (control and *Ltf*^*iCre*/+^*Arid1a*^*f/f*^ #1, n=6 IS; *Ltf*^*iCre*/+^*Arid1a*^*f/f*^ #2, n=3 IS). Scale bars: 100 μm. *(L)* Semi-quantitative H-score of stromal EGR1 staining strength in control versus *Ltf*^*iCre*/+^*Arid1a*^*f/f*^ #1. The graph shows the mean ± SEM (n=6 IS; *, p<0.05).

To determine if pSTAT3-EGR1 pathway dysregulation contributes to the implantation and decidualization defects of *Ltf*^*iCre*/+^*Arid1a*^*f/f*^ mice, we continued IHC analysis through the implantation window. While pSTAT3 and EGR1 were strongly expressed in decidualizing cells of control mice at GD 4.5, these proteins were significantly reduced in *Ltf*^*iCre*/+^*Arid1a*^*f/f*^ mice in the stroma surrounding embryos (−8.86 fold, p<0.0001; −1.66 fold, p=0.0002; respectively; Fig. 6*E-H*). This finding combined with the known importance of pSTAT3 and EGR1 in decidualization (38, 40) implicates dysregulation of the pSTAT3-EGR1 pathway in the decidualization defect of *Ltf*^*iCre*/+^*Arid1a*^*f/f*^ mice at GD 4.5. At GD 5.5, pSTAT3 and EGR1 expression normally decrease in the decidua (38, 39). In *Ltf*^*iCre*/+^*Arid1a*^*f/f*^ uterine sections, the implantation sites matching the major phenotype previously identified by histology (*Ltf*^*iCre*/+^*Arid1a*^*f/f*^ #1) exhibited significantly increased pSTAT3 and EGR1 expression in the stroma compared to controls (3.72 fold, p=0.0002; 3.24 fold, p=0.0411; respectively), whereas the minority phenotype (*Ltf*^*iCre*/+^*Arid1a*^*f/f*^ #2) was apparently unchanged (Fig. 6*I-L*). These findings reveal that the endometrial pSTAT3 and EGR1 activity critical for successful implantation is spatiotemporally dysregulated throughout early pregnancy as a result of epithelial *Arid1a* deletion.

Implantation and decidualization failure in *Pgr*^*cre*/+^*Foxa2*^*f/f*^ mice can be rescued by LIF repletion (10, 13, 32). To determine if the implantation defect in *Ltf*^*cre*/+^*Arid1a*^*f/f*^ mice was due primarily to decreased LIF expression at preimplantation and failure to activate the pSTAT3-EGR1 pathway, we administered recombinant LIF or vehicle (saline) at GD 3.5 and analyzed the uteri at GD 5.5. In both the vehicle and LIF-treated *Ltf*^*cre*/+^*Arid1a*^*f/f*^ mice, no successful implantation sites were evident morphologically or histologically (Fig. S5*A-B*). Additionally, IHC revealed strong pSTAT3 and EGR1 expression around the embryos matching the majority phenotype based on histology (*Ltf*^*cre*/+^*Arid1a*^*f/f*^ #1) in both groups, consistent with our findings in untreated *Ltf*^*cre*/+^*Arid1a*^*f/f*^ mice and contrasting with control mice at this stage (Fig. S5*C-D*; Fig. 6*I, K*).

### FOXA2 and ARID1A are Attenuated in Tandem in Women and Non-Human Primates with Endometriosis

To determine if ARID1A attenuation in women with endometriosis compromises endometrial gland function as found in mice, we examined FOXA2 expression in eutopic endometrial samples from women with and without endometriosis using IHC. Consistent with previous findings (15, 18), FOXA2 expression was strong across the secretory and proliferative phases of the menstrual cycle in the endometrial glands of control women and women with endometriosis (Fig. 7*A-B*). However, FOXA2 was decreased in women with endometriosis, and this decrease reached a level of statistical significance in secretory phase endometrium based on IHC H-score of staining strength in endometrial glands (−1.97 fold, p<0.01; Fig. 7*A-B*). Moreover, correlation analysis of FOXA2 and ARID1A gland-specific H-scores in serial secretory phase endometrial sections from women with endometriosis revealed a significant correlation between FOXA2 and ARID1A expression (Spearman correlation coefficient r=0.5982, p=0.0308; Fig. 7*C-D*). This finding supports the translational relevance of ARID1A’s role in endometrial gland function to human endometriosis.

**Figure 7.**
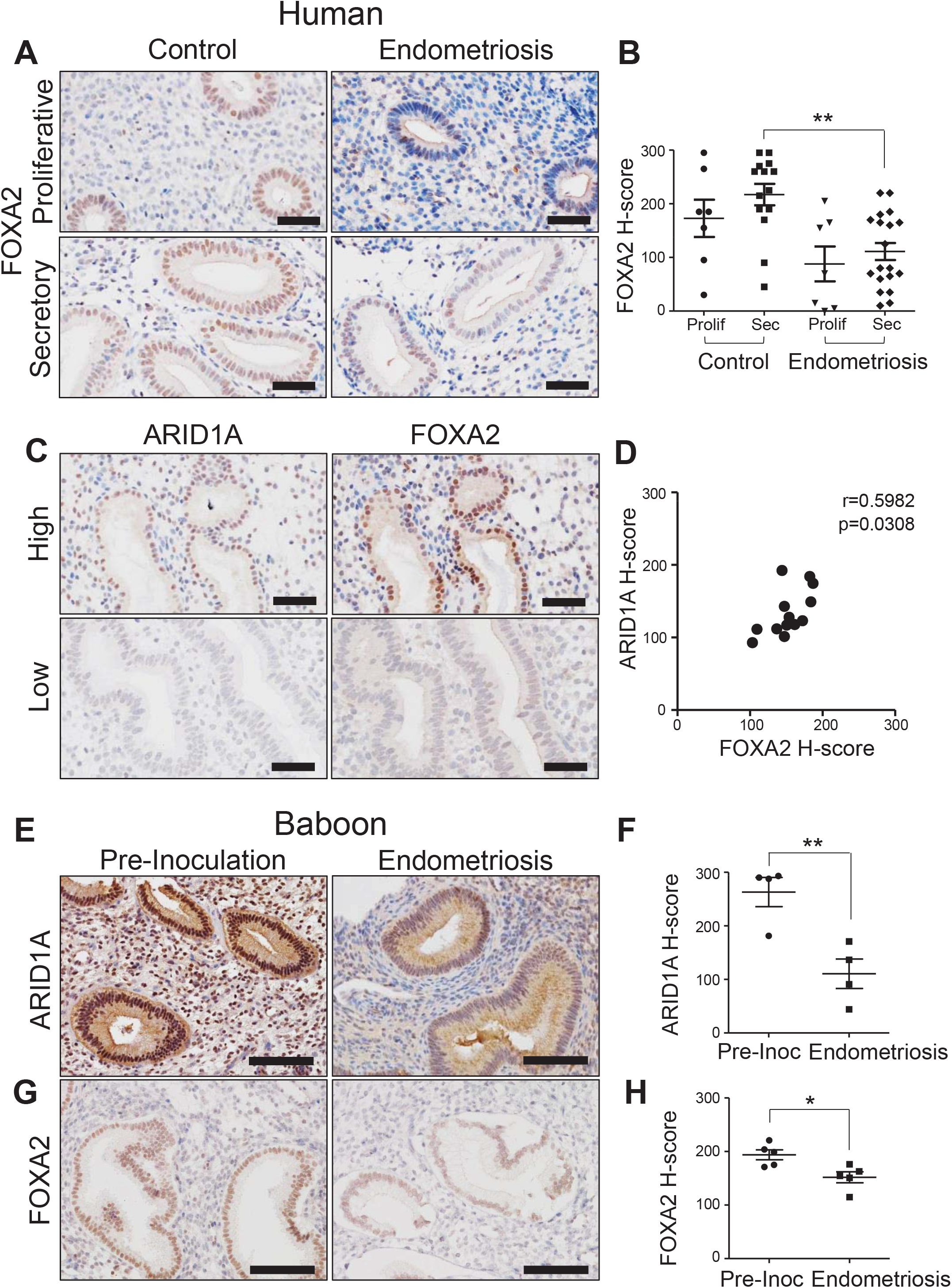
FOXA2 and ARID1A are Attenuated in Tandem in Women and Non-Human Primates with Endometriosis. *(A)* Representative images of FOXA2 IHC in endometrial biopsy samples from control women without endometriosis and women with confirmed endometriosis from the proliferative (n=7) and secretory (control n=14, endometriosis n=19) phases of the menstrual cycle. Scale bars: 50 μm. *(B)* Semi-quantitative H-score of FOXA2 staining strength in endometrial glands for the sample set pictured representatively in part *(A)*. The graph shows the mean ± SEM (**, p<0.01). *(C)* Representative images of strong and weak ARID1A and FOXA2 IHC in serial endometrial biopsy sections from women with endometriosis (n=13). Scale bars: 50 μm. *(D)* Correlation analysis of H-scores of FOXA2 staining strength and ARID1A staining strength in endometrial glands from IHC analysis of serial endometrial sections from women with endometriosis (n=13; p<0.0308). *(E)* Representative images of ARID1A IHC in paired endometrial biopsy samples from baboons before induction of endometriosis and after 1516 months of endometriosis development (n=4). Scale bars: 100 μm. *(F)* Semi-quantitative H-score of ARID1A staining strength in endometrial glands for the sample set pictured representatively in part *(E)*. The graph shows the mean ± SEM (n=4; **, p<0.01). *(G)* Representative images of FOXA2 IHC in paired endometrial biopsy samples from baboons before induction of endometriosis and after 1516 months of endometriosis development (n=5). Scale bars: 100 μm. *(H)* Semi-quantitative H-score of FOXA2 staining strength in endometrial glands for the sample set pictured representatively in part *(G)*. The graph shows the mean ± SEM (n=5; *, p<0.05).

To determine if endometriosis development alone is sufficient to cause reduction of ARID1A and FOXA2 in the eutopic endometrium, we utilized endometrial samples from a non-human primate model of endometriosis where menstrual effluent is inoculated into the peritoneal cavity to establish endometriotic lesions (26). Paired IHC analysis of samples taken before induction of endometriosis and 15-16 months after induction revealed significant decreases in both ARID1A (−2.38 fold, p=0.0034) and FOXA2 (−1.28 fold, p=0.0147) in baboon eutopic endometrial glands (Fig. 7*E-H*). These findings show that simply inducing development of endometriotic lesions is sufficient to reduce both ARID1A and FOXA2 levels in baboon eutopic endometrial glands.

## Discussion

ARID1A is primarily known for its roles in embryonic development (29) and as a tumor suppressor in many cancers, particularly endometriosis-related cancers of the female reproductive tract (19, 22, 41). However, recent findings from our group and others have established a role for ARID1A in normal uterine function and in cases of endometriosis without cancer (20, 21, 41, 42). In this study, we found that ARID1A plays an important role in endometrial gland development and function. We utilized three distinct mammalian systems to examine the importance and mechanism of ARID1A function in the endometrium: 1) conditional knockout mice revealed the temporal and compartment-specific physiological and molecular effects of *Arid1a* deletion in uterus; 2) endometrial biopsy samples from women with endometriosis confirmed the association between human endometriosis pathophysiology, decreased ARID1A expression, and endometrial gland dysfunction; and 3) a baboon model of endometriosis established a direct cause-effect relationship between endometriosis progression, ARID1A attenuation, and endometrial gland dysfunction.

Deletion of uterine *Arid1a* in mice (*Pgr*^*cre*/+^*Arid1a*^*f/f*^) compromised the ability of the endometrial epithelium to form typical numbers of glands by abrogating the expression of FOXA2, and our comparative analysis of transcriptomic data (20, 31) confirmed the molecular impact of uterine *Arid1a* loss on FOXA2-regulated genes. In particular, we found *Lif* expression diminished in the preimplantation *Pgr*^*cre*/+^*Arid1a*^*f/f*^ uterus, which is the key molecular dysfunction caused by *Foxa2* deletion resulting in implantation and decidualization defects (10, 13, 32). Though IHC results indicated *Pgr*^*cre*/+^*Arid1a*^*f/f*^ glands were FOXA2-negative, our method of analyzing uterine structure in 3D overcame the limitations of analyzing thin tissue sections to more holistically demonstrate a generalized failure of *Pgr*^*cre*/+^*Arid1a*^*f/f*^ uteri to develop the FOXA2-positive gland structures necessary to support early pregnancy establishment, in spite of a few scattered FOXA2-positive regions (9). Our ChIP-qPCR data from wildtype mice further revealed that uterine ARID1A regulation of FOXA2 expression during early pregnancy occurs in conjunction with direct binding of ARID1A at the *Foxa2* promoter.

Because of the defect of gland development resulting from early postnatal deletion of *Arid1a* in *Pgr*^*cre*/+^*Arid1a*^*f/f*^ mice, these mice are limited as a model of the ARID1A and FOXA2 downregulation in endometria of women with endometriosis that still appear to retain normal numbers of glands. Additionally, *Arid1a* is deleted in all uterine compartments of *Pgr*^*cre*/+^*Arid1a*^*f/f*^ mice (20, 27), which does not allow distinction between the epithelial and stromal functions of ARID1A. To overcome these limitations, we utilized *Ltf*^*iCre*/+^*Arid1a*^*f/f*^ mice to determine the effect of *Arid1a* deletion in the adult endometrial epithelium on reproductive function. *Ltf*^*iCre*/+^ mice do not express iCre until sexual maturity, and uterine expression is limited to the luminal and glandular epithethelium (28). However, *Ltf*^*iCre*/+^ mice also express iCre in some myeloid lineage immune cells (28, 43), so we cannot rule out the possibility of phenotypic effects resulting from *Arid1a* deletion in these cell types.

Our comprehensive characterization of the early pregnancy stage phenotypes of *Ltf*^*iCre*/+^*Arid1a*^*f/f*^ mice lends understanding of the epithelial-specific role of ARID1A in the endometrium, particularly in comparison with *Pgr*^*cre*/+^*Arid1a*^*f/f*^ mice (Table 1). Our data show that *Ltf*^*iCre*/+^*Arid1a*^*f/f*^ mice are severely sub-fertile due to uterine defects similar to those of *Pgr*^*cre*/+^*Arid1a*^*f/f*^ mice. However, though *Pgr*^*cre*/+^*Arid1a*^*f/f*^ mice exhibit major dysregulation of epithelial PGR signaling during early pregnancy (20), we found that this was not a major phenotype of *Ltf*^*iCre*/+^*Arid1a*^*f/f*^ mice, implying an important role for stromal ARID1A in coordinating epithelial PGR signaling. On the other hand, both conditional knockout models led to increased preimplantation epithelial proliferation, implying that suppression of this phenotype may be mediated in an epithelial cell-autonomous manner by ARID1A rather than through effects on stromal-epithelial juxtacrine signaling. Unlike *Pgr*^*cre*/+^*Arid1a*^*f/f*^ mice, *Ltf*^*iCre*/+^*Arid1a*^*f/f*^ mice had normal numbers of endometrial glands during early pregnancy, but *Ltf*^*iCre*/+^*Arid1a*^*f/f*^ mice still exhibited reductions of uterine *Foxa2* and *Lif* expression, which demonstrates normal gland structure formation but compromised gland function.

**Table 1.** Phenotype comparison of *Pgr*^*cre*/+^*Arid1a*^*f/f*^ and *Ltf*^*iCre*/+^*Arid1a*^*f/f*^ mice.

| Phenotype | *Pgr*<sup>Cre/+</sup>*Arid1a*<sup>ff</sup> | *Ltf*<sup>iCre/+</sup>*Arid1a*<sup>ff</sup> |
| --- | --- | --- |
| Fertility | Infertile (20) | Severely sub-fertile |
| Ovarian Function | Normal (20) | Normal |
| Implantation | Defect (open lumen) (20) | Defect (closed lumen) |
| Decidualization | Defect (20) | Defect |
| Uterine Receptivity | Non-receptive (20) | Non-receptive |
| Uterine Epithelial PGR Expression | Major decrease (20) | Minor decrease |
| Uterine Epithelial pESR1 Expression | Increased (20) | Normal |
| Endometrial Gland Number | Decreased | Normal |
| Uterine FOXA2 Expression | Decreased | Decreased |
| Uterine *Lif* Expression | Decreased | Decreased |

After LIF is secreted from endometrial glands at GD 3.5 and localizes to the glandular epithelium and sub-luminal stroma around the implanting embryo by GD 4.5 (44), it binds its transmembrane receptor and activates STAT3 via phosphorylation, then pSTAT3 translocates to the nucleus to induce EGR1 expression, which is rapid and transient in the sub-luminal stroma around the implanting embryo at this stage (39). Each step in this pathway is critical for implantation and decidualization success (33, 38, 40). Here, we show that *Ltf*^*iCre*/+^*Arid1a*^*f/f*^ mice fail to express normal amounts of *Lif* at GD 3.5 and experience disrupted pSTAT3 and EGR1 expression before, during, and after the implantation period. Notably, *Egr1* knockout mice exhibit increased epithelial proliferation, decreased stromal proliferation, and compromised decidualization (40), matching our finding in *Ltf*^*iCre*/+^*Arid1a*^*f/f*^ mice and providing a potential molecular explanation for the non-receptive endometrium phenotype. A failure to secrete sufficient amounts of LIF from ARID1A-deficient glands to activate STAT3 signaling in the stroma could thus explain how an epithelial cell defect of ARID1A compromises decidualization, a stromal cell process (Table 2).

**Table 2.** Expression patterns of ARID1A and ARID1A-regulated molecules in the peri-implantation mouse endometrium.

| **Molecule** | **Endometrial Expression** | **Cellular**<br>**Localization** | <sup>667</sup><br>**Reference** |
| --- | --- | --- | --- |
| ARID1A | Epithelium and stroma | Nucleus | (20) |
| FOXA2 | Glandular epithelium | Nucleus | (10) |
| LIF | Glandular epithelium and sub-luminal stroma around embryo | Secreted | (44) |
| pSTAT3 | Epithelium and stroma | Nucleus | (38) |
| EGR1 | Sub-luminal stroma around embryo | Nucleus | (39) |

The varying phenotypes found in *Ltf*^*iCre*/+^*Arid1a*^*f/f*^ mice at GD 5.5 appear to represent a failure of the decidualization process at two different stages: one at the stage characteristic of the normal GD 4.5 expression patterns of pSTAT3 and EGR1 (*Ltf*^*iCre*/+^*Arid1a*^*f/f*^ #1), and one matching the normal GD 3.5 patterns *Ltf*^*iCre*/+^*Arid1a*^*f/f*^ #2). The source of the variation in these results is unclear, but it could possibly have resulted from slightly different timing from mating event to sample collection between mice or from localized variation in iCre activity.

Overall, our data from *Ltf*^*iCre*/+^*Arid1a*^*f/f*^ mice support the hypothesis that epithelial ARID1A is necessary to potentiate the FOXA2 expression and tightly regulated LIF-STAT3-EGR1 pathway signaling required in the uterus to support implantation and decidualization. However, restoring LIF levels alone did not reverse the implantation failure that resulted from loss of epithelial ARID1A, which implies that deletion of *Arid1a* in the adult endometrial epithelium must cause other molecular defects besides reduction of FOXA2 expression that preclude early pregnancy establishment.

Genetically engineered mice are powerful tools to study the molecular regulation of early pregnancy due to the similarity between mouse and human reproduction (3); however, studies in higher primates and directly in women are necessary to draw stronger conclusions about the relevance of any findings to human pathophysiology. We previously found a reduction of ARID1A expression in the endometrium of infertile women with endometriosis (20). Here, our findings were consistent with reports that FOXA2 expression is markedly reduced in endometrium from women with endometriosis (15, 16), and we demonstrated a correlation between endometrial ARID1A and FOXA2 expression among women with endometriosis. This finding supports the idea that ARID1A regulates FOXA2 and gland function in women as well as in mice and that this function is compromised by endometriosis. However, no direct cause-effect relationship between endometriosis and ARID1A-FOXA2 expression can be established with this type of associational study: it is not clear whether endometriosis causes this particular molecular dysfunction or if it is a risk factor for endometriosis development. The use of a non-human primate model of induced endometriosis allows paired sample analysis before and after disease development in a menstruating species biologically similar to humans (26). In this way, our study in baboons establishes that the presence of endometriotic lesions precipitates the parallel downregulation of ARID1A and FOXA2 in the eutopic endometrium, indicating that endometriosis pathophysiology causes disruption of ARID1A’s regulation of endometrial gland function.

Worthy of note, a recent report showed that endometrial epithelial expression of the transcription factor sex-determining region Y-related high-mobility group box protein 17 (SOX17) is critical for successful implantation and decidualization in mice and is reduced in the endometrial glands of women with endometriosis (45). Interestingly, transcriptomic analysis of endometrial epithelial specific SOX17 knockout mice revealed that SOX17-regulated gene expression patterns remarkably overlapped with ARID1A-regulated genes and FOXA2-regulated genes (45). Furthermore, ablation of SOX17 diminished the protein levels of ARID1A and FOXA2 in the endometrium, which when taken together with our data, indicates a possible hierarchical relationship between these three proteins in the endometrium, with SOX17 positively regulating ARID1A which in turn promotes FOXA2 expression and gland function (45).

Our complementary experimental methods utilizing samples from mice, women, and non-human primates coordinate to reveal the critical function of epithelial ARID1A in the endometrium that is compromised in cases affected by endometriosis. As more is determined about the connection between endometriosis and infertility, ARID1A will continue to be a crucial molecular factor to consider when developing potential therapies to combat the effects of these common and devastating conditions.

## Acknowledgments

Research reported in this publication was supported in part by the Eunice Kennedy Shriver National Institute of Child Health & Human Development of the National Institutes of Health under Award Number R01HD084478 to J.W.J., F31HD101207 and T32HD087166 to R.M.M., and P50HD28934 (NCTRI) to The University of Virginia Center for Research in Reproduction Ligand Assay and Analysis Core; MSU AgBio Research, and Michigan State University. The content is solely the responsibility of the authors and does not necessarily represent the official views of the National Institutes of Health. We are grateful for the generous donation of *Arid1a*^*f/f*^ mice by Dr. Zhong Wang, University of Michigan. We would like to thank Erin Vegter and Samantha Hrbek for technical support.

## Author Contributions

R.M.M., T.H.K., R.A., and J.W.J designed the experiments and analyzed data; R.M.M., J.Y.Y., T.H.K., H.E.T., and R.A. performed experiments; A.T.F., S.L.Y., and B.A.L. collected the endometrial samples and interpreted the results; R.M.M., T.H.K., and J.W.J wrote the manuscript.

---

**Figure S1.**
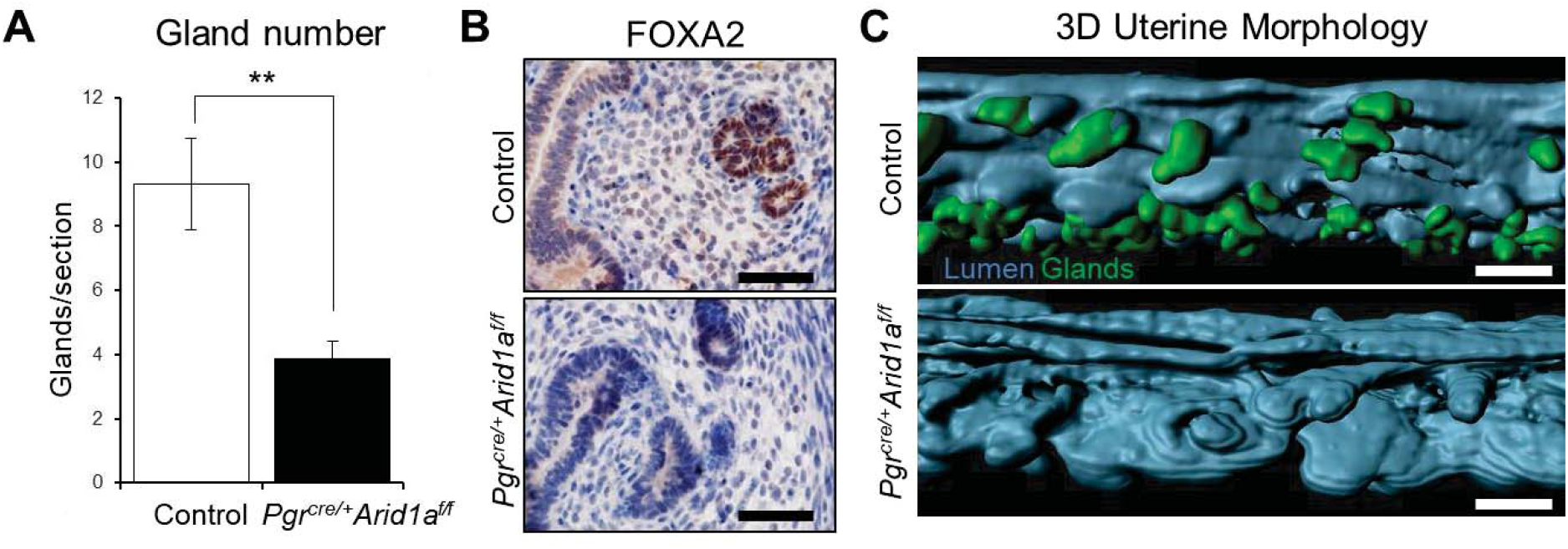
Endometrial glands are dysregulated during postnatal development in *Pgr*^*cre*/+^*Arid1a*^*f/f*^ mice. *(A)* Endometrial gland counts in control and *Pgr*^*cre*/+^*Arid1a*^*f/f*^ mice at 4 weeks of age. The graph represents the mean ± SEM of the number of glands per uterine tissue section (n=8; **, p=0.01). *(B)* Representative images of FOXA2 IHC in control and *Pgr*^*cre*/+^*Arid1a*^*f/f*^ mouse uterine sections at 4 weeks of age (n=3). Scale bars: 50 μm. *(C)* Three-dimensional uterine morphology of control and *Pgr*^*cre*/+^*Arid1a*^*f/f*^ uterine horns at 3 weeks of age based on whole-mount immunofluorescence for E-cadherin and FOXA2, where the 3D luminal structure (blue) is constructed by subtracting the FOXA2 (green) from the E-cadherin signal (Control, n=3; *Pgr*^*cre*/+^*Arid1a*^*f/f*^, n=2). Scale bars: 100 μm.

**Figure S2.**
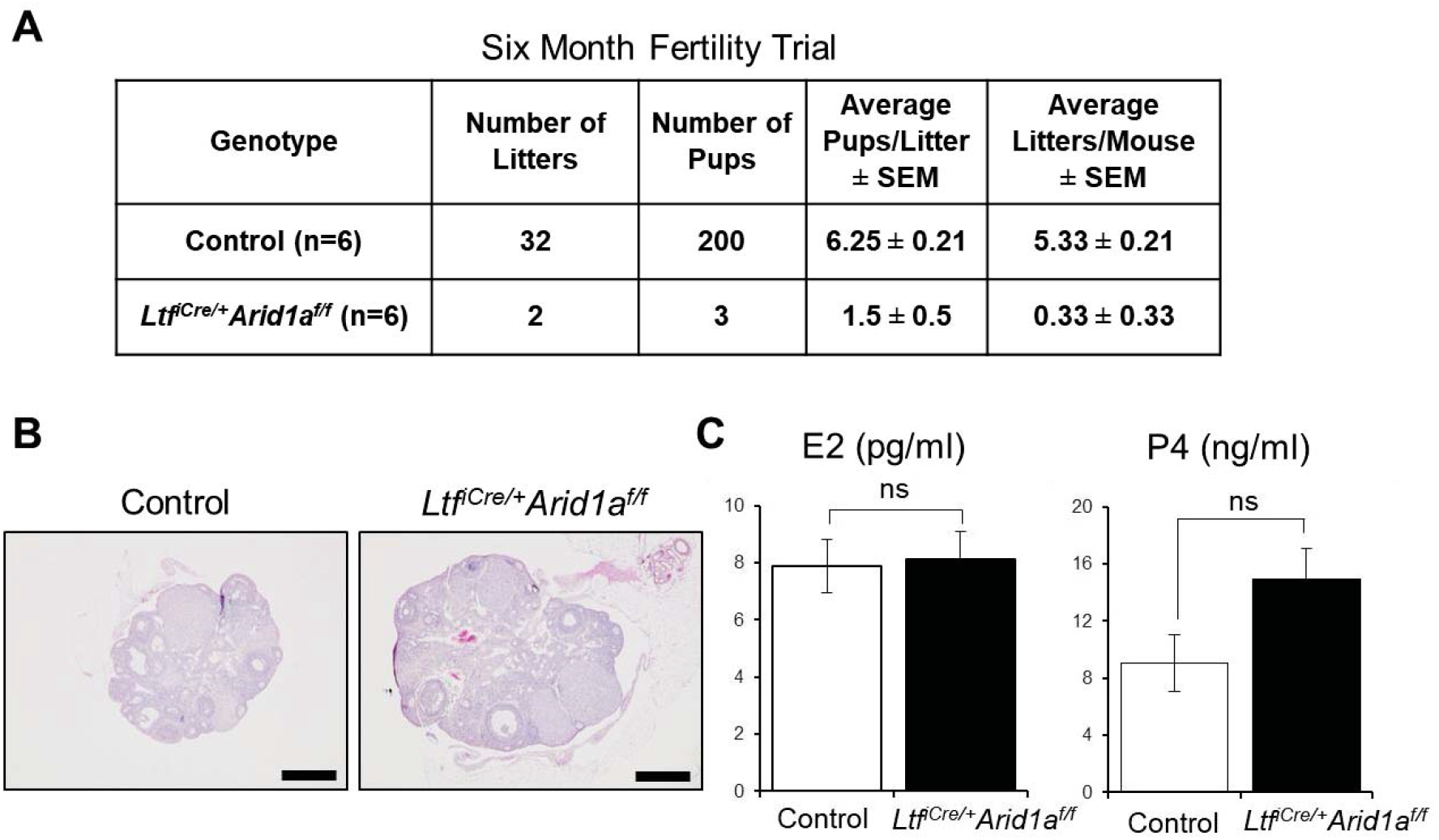
Endometrial Epithelial-Specific *Arid1a* Loss Results in Severe Sub-Fertility but Normal Ovarian Function. *(A)* Resulting numbers of litters and pups found in a fertility trial where female control or *Ltf*^*iCre*/+^*Arid1a*^*f/f*^ mice were housed in breeding cages with wildtype male mice for six months. *(B)* Representative images of hematoxylin and eosin-stained ovary cross-sections from control and *Ltf*^*iCre*/+^*Arid1a*^*f/f*^ mice at GD 3.5 (n=3). Scale bars: 500 μm. *(C)* Quantification of serum P4 and E2 levels in control and *Ltf*^*iCre*/+^*Arid1a*^*f/f*^ mice at GD 3.5. The graphs represent the mean ± SEM (n=5; ns, p>0.05).

**Figure S3.**
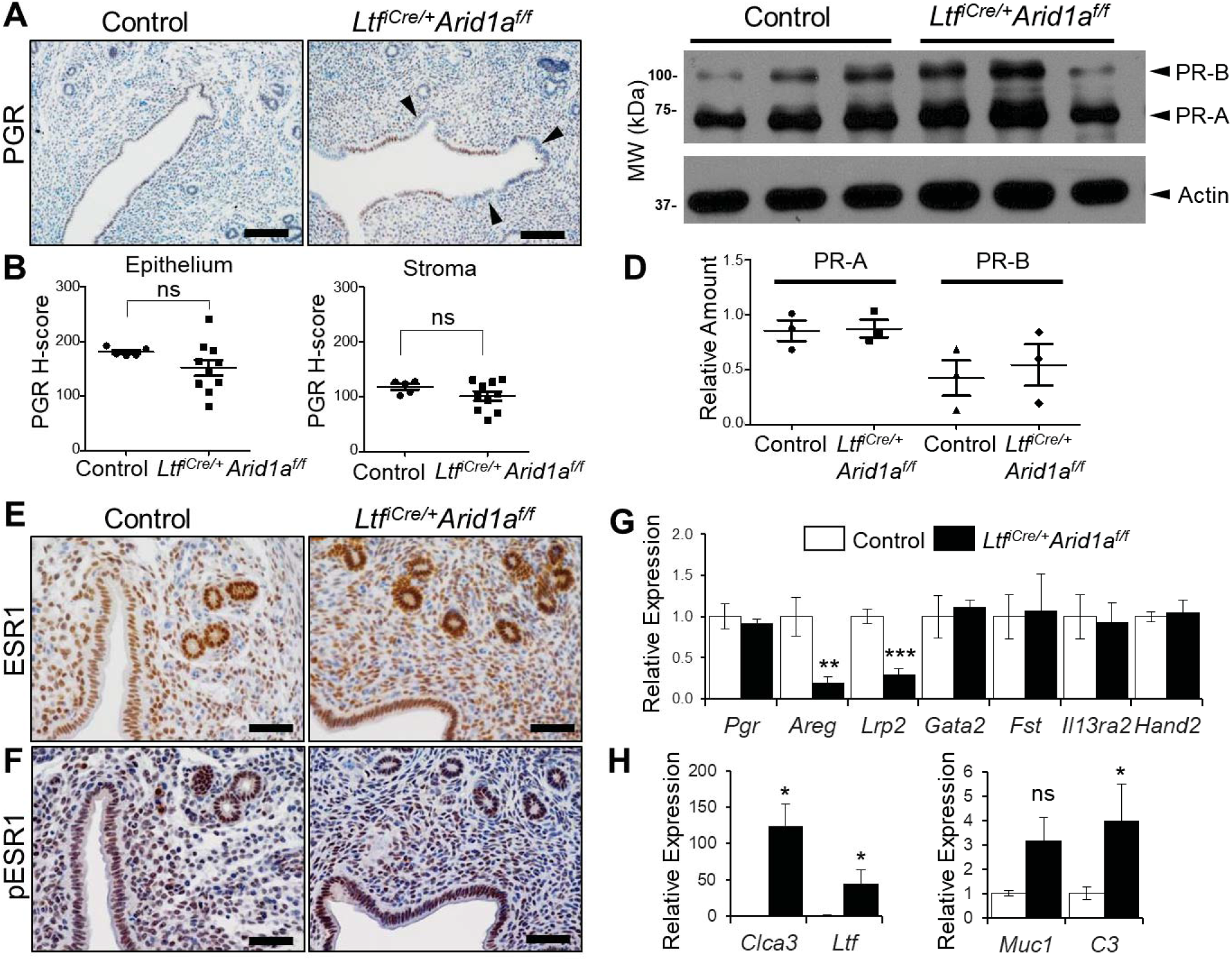
P4 and E2 Signaling Exhibit Minor Changes in GD 3.5 *Ltf*^*iCre*/+^*Arid1a*^*f/f*^ Mice. *(A)* Representative images of PGR IHC in control and *Ltf*^*iCre*/+^*Arid1a*^*f/f*^ mouse uterine tissue sections (control, n=3; *Ltf*^*iCre*/+^*Arid1a*^*f/f*^, n=10). Arrowheads indicate patches of PGR-negative cells in the luminal epithelium. Scale bars: 100 μm. *(B)* Semi-quantitative H-scores of epithelial and stromal PGR staining strength. The graphs represent the mean ± SEM (control, n=3, 5 tissue regions; *Ltf*^*iCre*/+^*Arid1a*^*f/f*^, n=10; ns, p>0.05). *(C)* Western blot of PGR (PR-A and PR-B) in protein isolated from total uterine tissue of control and *Ltf*^*iCre*/+^*Arid1a*^*f/f*^ mouse uteri (n=3). Western blotting was performed at previously described (1). *(D)* Quantification of band intensity of PR-A and PR-B relative to actin in control and *Ltf*^*iCre*/+^*Arid1a*^*f/f*^ uteri (n=3, p>0.05). *(E)* Representative images of ESR1 IHC in control and *Ltf*^*iCre*/+^*Arid1a*^*f/f*^ mouse uterine tissue sections (n=3). Scale bars: 50 μm. *(F)* Representative images of pESR1 IHC in control and *Ltf*^*iCre*/+^*Arid1a*^*f/f*^ mouse uterine tissue sections (n=3). Scale bars: 50 μm. *(G)* Relative expression of P4 target gene mRNA normalized to *Rpl7* in whole uterine tissue preparations. The graphs represent the mean ± SEM (control, n=4; *Ltf*^*iCre*/+^*Arid1a*^*f/f*^, n=5; **, p<0.01; ***, p<0.001). *(H)* Relative expression of E2 target gene mRNA normalized to *Rpl7* in whole uterine tissue preparations. The graphs represent the mean ± SEM (control, n=4; *Ltf*^*iCre*/+^*Arid1a*^*f/f*^, n=5; *, p<0.05).

**Figure S4.**
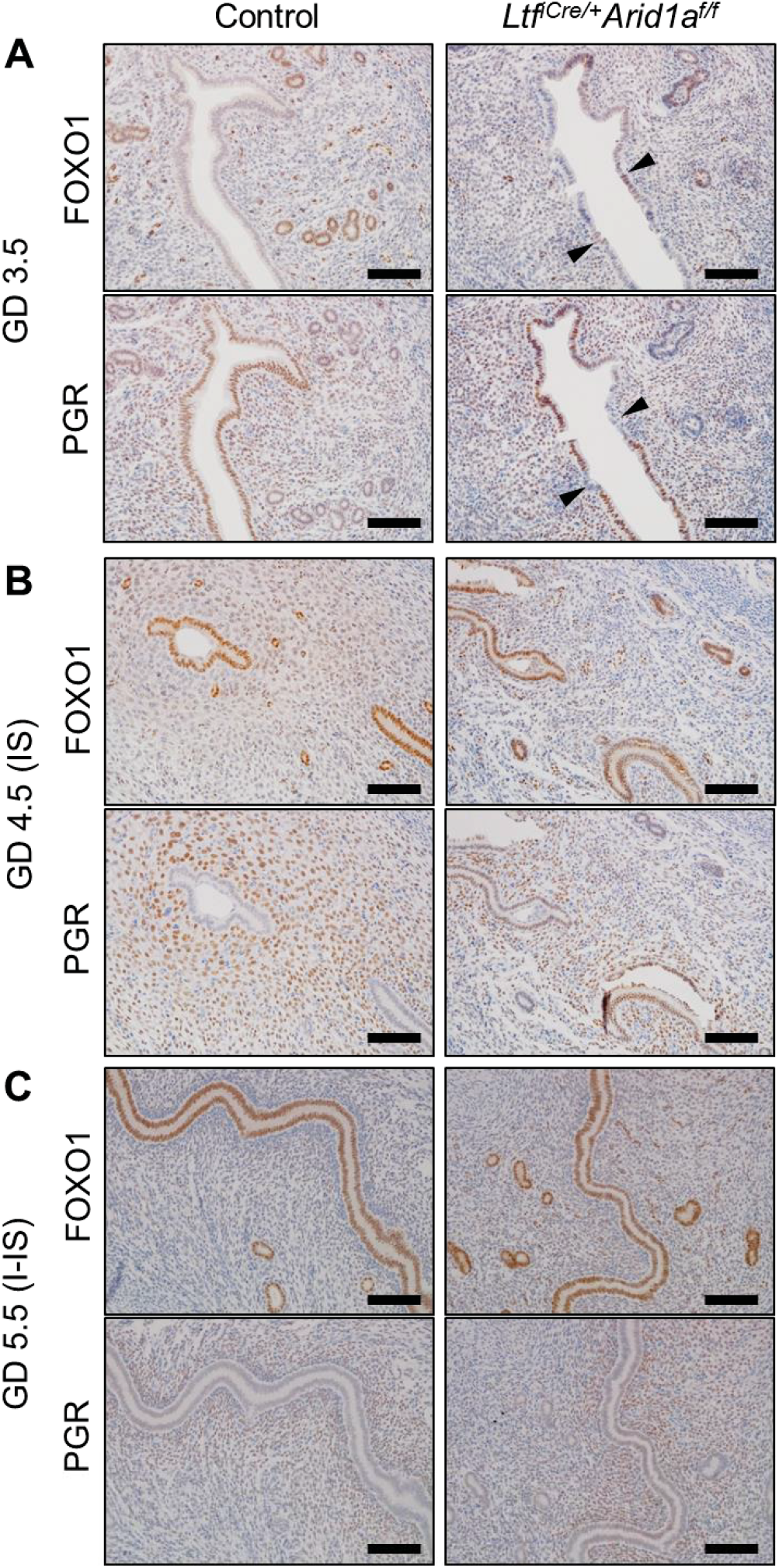
FOXO1 and PGR are Not Notably Altered in the *Ltf*^*iCre*/+^*Arid1a*^*f/f*^ Uterus during Early Pregnancy. *(A)* Representative images of FOXO1 (upper panel) and PGR (lower panel) IHC in control and *Ltf*^*iCre*/+^*Arid1a*^*f/f*^ mouse serial uterine tissue sections at GD 3.5 (n=3). Arrowheads indicate patches of FOXO1-positive/PGR-negative cells in the luminal epithelium. Scale bars: 100 μm. *(B)* Representative images of FOXO1 (upper panel) and PGR (lower panel) IHC in control and *Ltf*^*iCre*/+^*Arid1a*^*f/f*^ mouse serial uterine tissue sections surrounding implantation sites (IS) at GD 4.5 (n=3, 6 IS). Scale bars: 100 μm. *(C)* Representative images of FOXO1 (upper panel) and PGR (lower panel) IHC in control and *Ltf*^*iCre*/+^*Arid1a*^*f/f*^ mouse serial uterine tissue sections at inter-implantation site regions (I-IS) at GD 5.5 (control, n=3; *Ltf*^*iCre*/+^*Arid1a*^*f/f*^, n=5). Scale bars: 100 μm.

**Figure S5.**
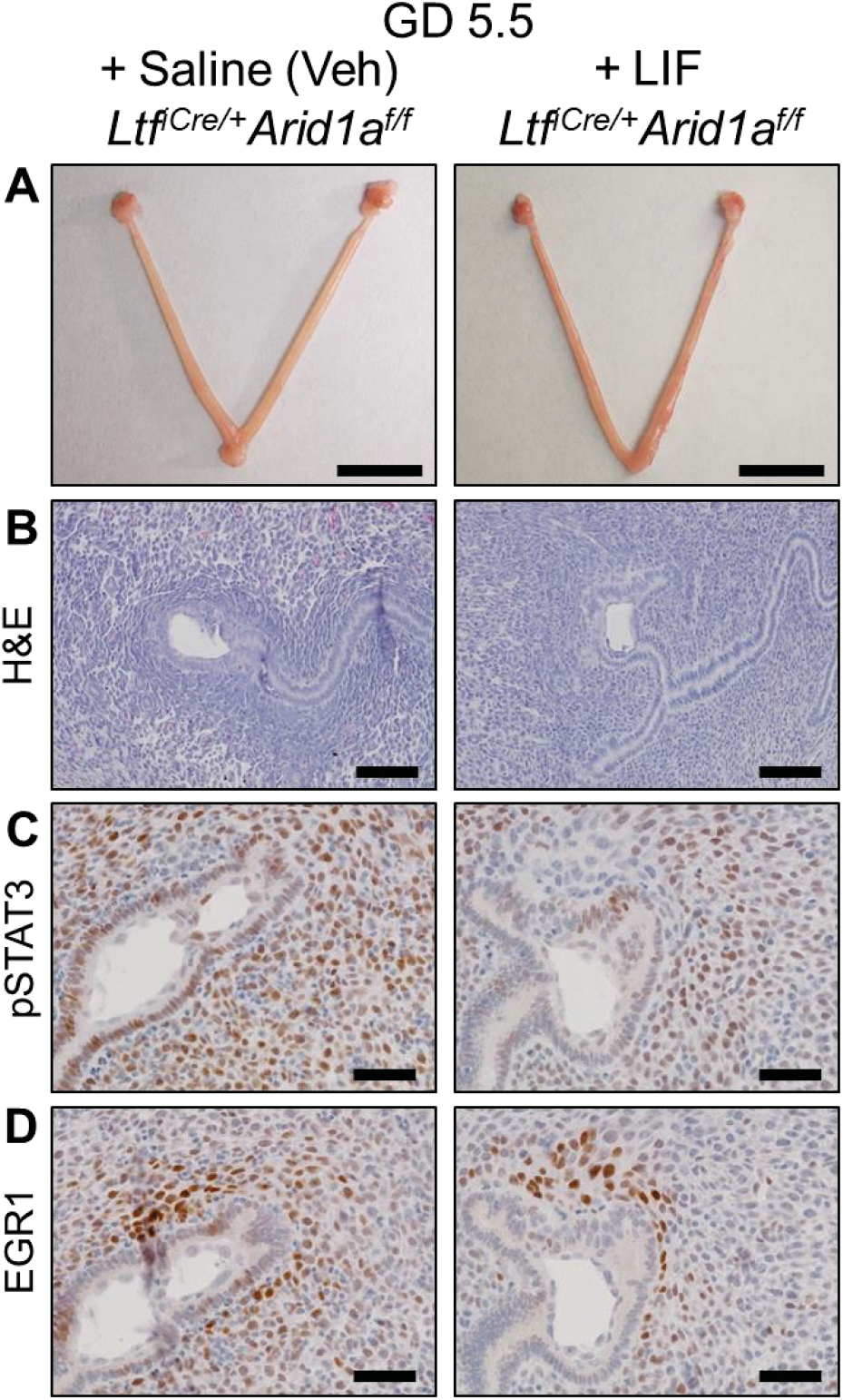
LIF Repletion at GD 3.5 Does Not Rescue Implantation in *Ltf*^*iCre*/+^*Arid1a*^*f/f*^ Mice. *(A)* No implantation sites were grossly visible at GD 5.5 in uteri from vehicle (Veh) or LIF-treated *Ltf*^*iCre*/+^*Arid1a*^*f/f*^ mice (n=3). Scale bars: 1 cm. Attempted rescue of implantation by LIF repletion was performed as described previously (2). Briefly, *Ltf*^*iCre*/+^*Arid1a*^*f/f*^ mice received i.p. injections of 10 μg recombinant mouse LIF (BioLegend, San Diego, CA) in saline or vehicle (saline only) on GD 3.5 at 1000 and 1800 hours. Implantation sites were analyzed on the morning of GD 5.5 morphologically and histologically. *(B)* Representative images of hematoxylin and eosin-stained uterine tissue sections from control and *Ltf*^*iCre*/+^*Arid1a*^*f/f*^ mice at GD 5.5 (Veh-treated n=3, 7 IS; LIF-treated n=3, 9 IS) Scale bars: 100 μm. *(C)* Representative images of pSTAT3 IHC in saline (vehicle) and LIF-treated *Ltf*^*iCre*/+^*Arid1a*^*f/f*^ mouse uteri. Scale bars: 50 μm. *(D)* Representative images of EGR1 IHC in saline (vehicle) and LIF-treated *Ltf*^*iCre*/+^*Arid1a*^*f/f*^ mouse uteri. Scale bars: 50 μm.

**Table S1.** Antibody List.

| **Target** | **Company** | **Catalog #** |
| --- | --- | --- |
| ARID1A (IF) | Abnova | H00008289-M02 |
| ARID1A (IHC) | Santa Cruz | SC-98441 |
| ARID1A (ChIP) | Abnova | MAB15809 |
| COX-2 | Cayman | 160106 |
| E-cadherin (IHC) | Cell Signaling | 3195 |
| E-cadherin (3D, IF) | Clontech | M108 |
| EGR1 | Cell Signaling | 4153 |
| FOXA2 (IHC) | Seven Hills | WRAB-FOXA2 |
| FOXA2 (3D, IF) | Novus Biologicals | NBP1-95426 |
| FOXO1 | Cell Signaling | 2880 |
| Ki67 | BD Pharmingen | 550609 |
| PGR | Cell Signaling | 8757 |
| ESR1 | Vector | VP-E613 |
| pESR1 | Abcam | ab31477 |
| STAT3 | Cell Signaling | 4904 |
| pSTAT3 | Cell Signaling | 9145 |

**Table S2.**
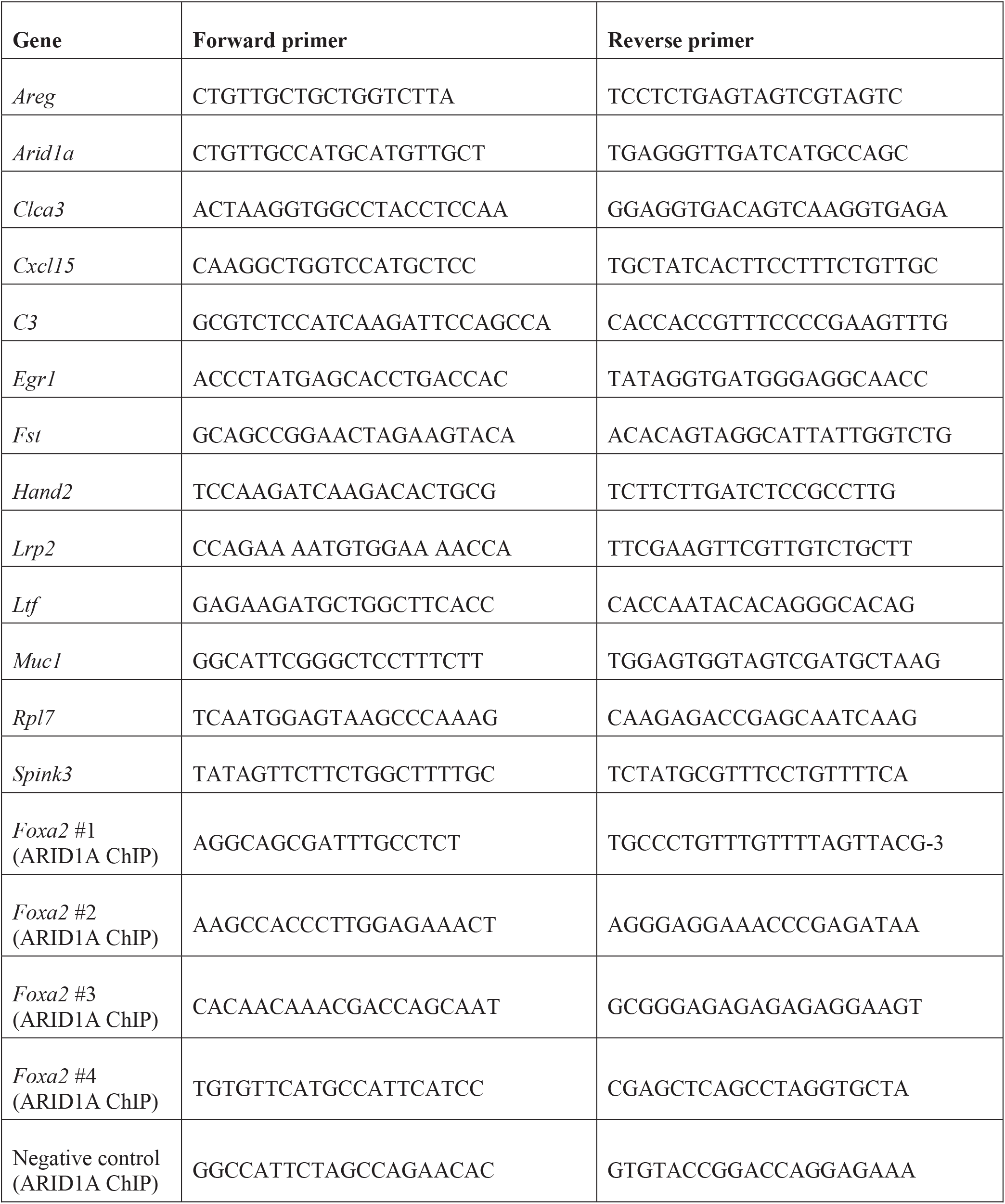
SYBR Green qPCR Primer List.

**Table S3.**
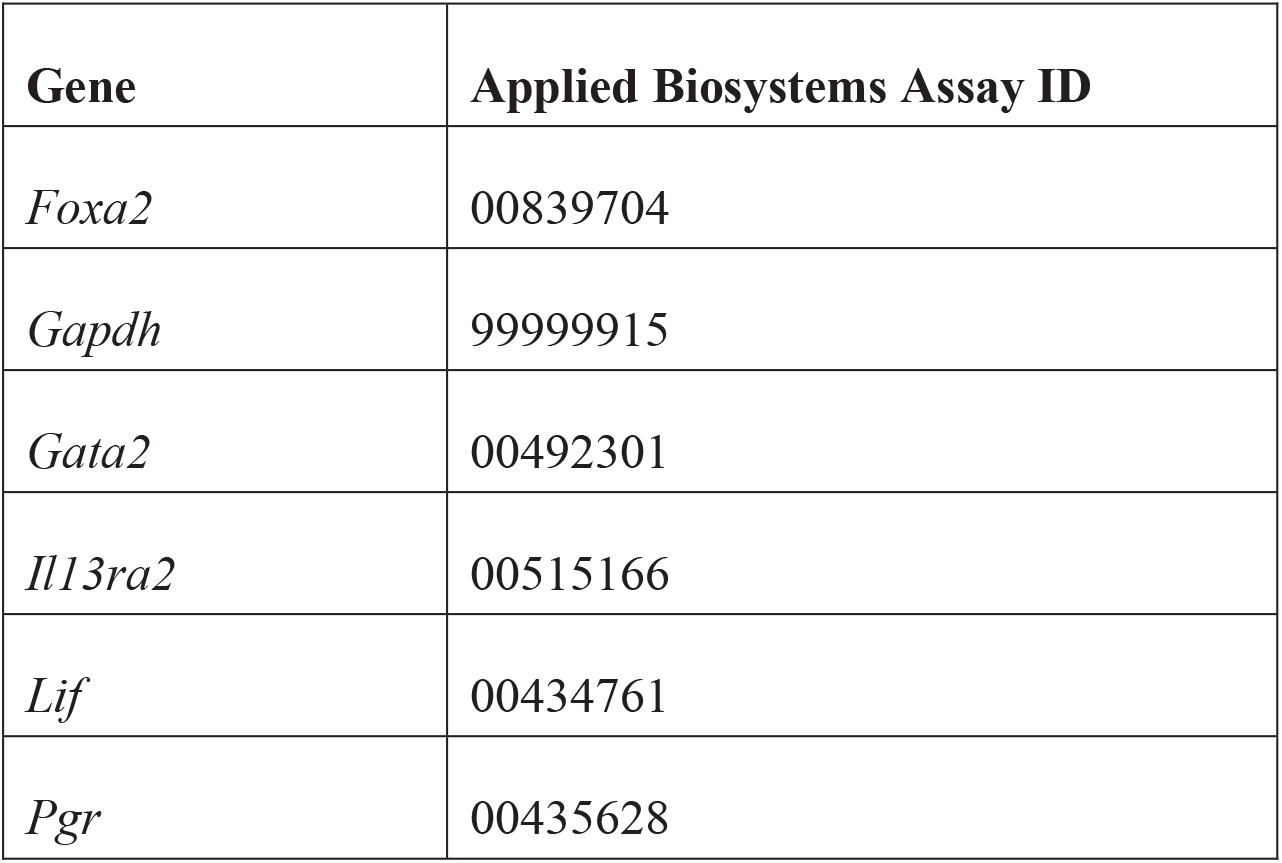
TaqMan qPCR probe list.

| **Gene** | **Applied Biosystems Assay ID** |
| --- | --- |
| *Foxa2* | 00839704 |
| *Gapdh* | 99999915 |
| *Gata2* | 00492301 |
| *Il13ra2* | 00515166 |
| *Lif* | 00434761 |
| *Pgr* | 00435628 |

**Dataset S1. Genes dysregulated in the *Pgr*^*cre*/+^*Arid1a*^*f/f*^ and *Pgr*^*cre*/+^*Foxa2*^*f/f*^ uterus at GD 3.5.**

The supplementary dataset includes gene identifiers, *Pgr*^*cre*/+^*Arid1a*^*f/f*^ fold change, and *Pgr*^*cre*/+^*Foxa2*^*f/f*^ fold change of genes differentially expressed in both the *Pgr*^*cre*/+^*Arid1a*^*f/f*^ and the *Pgr*^*cre*/+^*Foxa2*^*f/f*^ uterus at GD 3.5. As represented in Fig. 1D, 316 genes are dysregulated at GD 3.5 in the uterus by uterine deletion of *Arid1a* or *Foxa2*. To generate this list, we compared differentially expressed genes from published *Pgr*^*cre*/+^*Arid1a*^*f/f*^ transcriptomics analysis (3) with differentially expressed genes from published transcriptomics analysis of *Pgr*^*cre*/+^*Foxa2*^*f/f*^ mice (4). Duplicate genes were first removed based on GeneBank ID or gene symbol.

